# Aerial litter mimicry: a novel form of floral deception mediated by a monoterpene synthase

**DOI:** 10.1101/2024.06.12.596753

**Authors:** Ming-Fai Liu, Junhao Chen, Katherine R. Goodrich, Sung Kay Chiu, Chun-Chiu Pang, Tanya Scharaschkin, Richard M. K. Saunders

## Abstract

1. Floral mimics deceive their pollinators by developing visual and olfactory resemblance to their models. Our knowledge on the diversity of models is expanding rapidly. We report a system in which the flowers exhibit phenotypes similar to aerial litter and deceives an aerial litter specialist beetle to achieve pollination.
2. We assessed the floral phenology and the effective pollinators of an Australian understorey treelet, *Meiogyne heteropetala* (Annonaceae). The similarities of morphology, colour and odour between the flowers and co-occurring aerial litter were investigated. The terpene synthase involved in floral scent emission was identified by expression patterns and product profile. The behavioural responses of the pollinator to various odours were assessed using bioassays.
3. The erotylid beetle *Loberus sharpi* is the most likely effective pollinator of *M. heteropetala*, and its eggs were found on the petals of *M. heteropetala*. *Loberus sharpi* was exclusively found in aerial litter and *M. heteropetala* flowers. The morphology and spectral reflectance of the flowers overlap with aerial litter. The floral scent was dominated by monoterpenes, especially 1,8-cineole. The cineole synthase MhCINS was the only highly expressed floral terpene synthase and possessed a highly similar product profile to the floral scent composition. NMDS showed that the volatile composition of *M. heteropetala* flowers is distinct from other congeners and highly similar to aerial litter, indicating advergence to aerial litter. Visual and odour resemblance, coupled with the deposition of eggs on the flowers, provides evidence that the beetles were deceived into pollinating the flowers. Behavioural experiments showed that the pollinator was attracted to both aerial litter and *M. heteropetala* flowers. The beetles were also attracted to 1,8-cineole and synthetic mixes of floral odour and MhCINS products. The beetles were unable to distinguish floral scent from MhCINS products, nor from 1,8-cineole, suggesting MhCINS alone sufficed to attract the pollinator olfactorily. The beetles, however, preferred aerial litter over flowers. The beetles likely categorised the flower as a general, but not the most preferred brood substrate.
4. *Synthesis.* This study reports the first case of floral mimicry of aerial litter and characterises the biochemical process responsible for olfactory mimicry.

## Introduction

In floral mimicry systems, the selective force exerted by pollinators has led mimetic flowers to develop extraordinary resemblance to their respective models (Johnson & Schiestl, 2016; Gaskett, 2011). These plants evolved a diverse array of highly specialised floral secondary metabolites (Wong et al., 2017a, 2023) and unusual morphologies, prompting the pollinators into miscategorising the flowers as a variety of organisms and substrates, including oviposition sites, food sources, or conspecific mates (Jersáková, et al., 2009; Gaskett, 2011; Urru et al., 2011). This misclassification by the animal results in repeated floral visitation and thus pollination (Dafni, 1984; Roy & Widmer, 1999; Schaefer & Ruxton, 2009; Vereecken & McNeil, 2010; Schiestl & Johnson, 2013). These flowers display advergent evolution, with floral phenotypes unilaterally evolving towards the phenotype of the model, but not vice versa (Johnson & Schiestl, 2016). The most prevalent form of floral mimicry is arguably oviposition-site mimicry, which can be found in more than 20 families, including Annonaceae, Apocynaceae, Araceae, Aristolochiaceae, and Orchidaceae, (Urru et al., 2011; Jürgens et al., 2013; Jürgens & Shuttleworth, 2015). These flowers fool insects into visiting and often laying eggs in the flowers by imitating cues that insects use in search for their brood sites such as dung, carcasses, mushrooms and aphid-infested vegetation (Bänziger, 1996a; Jürgens & Shuttleworth, 2015; Policha et al., 2016). Animal by-products, especially dung and carcasses, are among the most widely utilised models (Urru et al., 2011; Jürgens et al., 2013; Jürgens & Shuttleworth, 2015). Because eggs laid in these flowers almost always end in inevitable death, insect broods rarely damaged the developing ovules (Johnson & Schiestl, 2016). Models of plant origin are rare in oviposition-site mimicry, and are largely confined to fruits (Jürgens et al., 2000; Goodrich & Jürgens, 2018).

Aerial litter (also known as canopy litter or arboreal litter) refers to the arboreal accumulation of dead plant material in the forest canopy and undergrowth. Aerial litter can arise from self-retention of dead tissues (Yoshida & Hijii, 2006; Matsushita et al., 2017), or interception of fallen litter by the lower canopy, undergrowth, and epiphytes (Nadkarni & Matelson, 1991; Zona & Christenhusz, 2015). Some plants and fungi develop specialized morphologies and structures to trap falling litter and utilise its nutrients (Hedger et al., 1993; Snaddon et al., 2012; Zona & Christenhusz, 2015). Quantities of leaf litter retained likely vary widely across habitat types based on wind velocity (Nadkarni & Matelson, 1991) and floristic composition (Zona & Christenhusz, 2015), with some plants capable of retaining or capturing *ca.* 10–50% incoming litter (Alvarez-Sanchez & Guevara, 1999; Dearden & Wardle, 2008). Arboreal environments display lower humidity and greater moisture fluctuation than the forest floor (Bohlman et al., 1995; Lindo & Winchester, 2007). They provide important microhabitats for various decomposers such as fungi, dictyostelids, myxomycetes and protostelids (Lodge & Cantrell, 1995; Stephenson et al., 1999; Black et al., 2004; Marques et al., 2015) and arthropods, including Arachnida, Blattodea, Coleoptera, Collembola, Hymenoptera and Orthoptera (Greenberg, 1987; Nadkarni & Longino, 1990; Rosenberg, 1993; Rosenberg, 1997; Yoshida & Hijii, 2005; Snaddon et al., 2012; Mansor et al., 2019). Many fungi, slime molds and arthropods are aerial litter specialist (Longino & Nadkarni, 1990; Moore & Stephenson, 2003; Fagon et al., 2006; Marques et al., 2015). The intricate relationship between aerial litter and insects entails established communication channels, which are exploitable by floral mimics.

The Annonaceae is an early-divergent pantropical family of trees and lianas with mostly ‘mess-and-soil’ pollination strategy (Faegri & van der Pijl, 1966; Pang & Saunders, 2014), in which flowers with primitive architecture are pollinated primarily by beetles in the families Curculionidae, Nitidulidae, Scarabaeidae, and Staphylinidae (Gottsberger, 1999; Saunders, 2012). These flowers serve as tryst sites, where beetles spend considerable time consuming pollen, stigmatic exudate and floral tissues as rewards (Gottsberger & Webber, 2018; Saunders, 2020). Although Annonaceae flowers are self-compatible, they are strongly protogynous and predominantly outcrossing, with no reports of obligate self-fertilisation (Pang & Saunders, 2014). Annonaceae have highly variable floral odour (Goodrich & Raguso, 2009; Goodrich, 2012), giving rise to a variety of pollination specialisations, such as euglossine bee pollination (Teichert et al., 2009), “private communication channel” with scarab beetles (Maia et al., 2021), and potential mimicry of fruits (Jürgens et al., 2000; Gottsberger, 2016; Chen et al., 2020), fermenting substrates (Goodrich et al., 2006; 2023) and mushrooms (Teichert et al., 2012). As nearly half of the species were described as having fruity floral odour (Goodrich, 2012), fruit mimicry might be prevalent and ancestral in the family (Gottsberger, 1999, 2016; Jürgens et al., 2000; Saunders, 2012; Goodrich & Jürgens, 2018). The Annonaceae genus *Meiogyne* likewise generally possesses a yellowish corolla with a fruity floral odour (Silberbauer-Gottsberger et al., 2003; Tan et al., 2014; Johnson et al., 2019; Xue et al., 2021). Some species however possess dark petals (Jessup, 2007). An example is the understorey treelet *Meiogyne heteropetala*, occurring in coastal and subcoastal notophyll forests (leaf area: 2,025–4,500 mm^2^; Webb et al., 1959) along the eastern coast of Queensland, Australia (Jessup, 2007). Although they were speculated to be sapromyophilous (i.e. emit foul scent to attract dung- or carrion-seeking flies; Saunders, 2020), pilot observations instead revealed that the dark maroon flowers produce a minty, bark-like scent and bear a striking resemblance to aerial litter in the vicinity. In this study, we address the following questions: (1) What are the most likely effective pollinators of *M. heteropetala*? (2) Does *M. heteropetala* display visual and olfactory advergence to aerial litter? (3) What is the biochemical pathway involved in scent production of *M. heteropetala*? (4) Does the enzyme play a role in pollinator attraction? By asking these questions, we aim to characterise the phenotypic basis on which *M. heteropetala* attract pollinators and elucidate the biochemical process for the hypothesised olfactory mimicry. This rarely adopted integrative approach would contribute significantly to our understanding of floral mimicry by uniting multiple lines of evidence.

### Materials and Methods Study species and sites

Field work was conducted during the peak flowering season of *Meiogyne heteropetala* (December–Feburary) in five years (2013, 2018, 2020, 2022, 2023) in the coastal notophyll forests of Bowling Green Bay National Park, Dingo Beach and Conway National Park in Queensland, Australia (Permit no.: WITK08297010 & WITK186669017-2). Each location comprises *ca.* 50 individuals. The field sites at Dingo Beach and Conway National Park are coastal closed forest, while the field site at Bowling Green Bay National Park is a eucalyptus semi-closed forest fringing the Bowling Green Bay Ramsar wetlands. The field sites are 35–127 km apart.

### Floral phenology

Floral ontogeny was studied in Dingo Beach, where 25 flowers from *ca.* 5 individuals were tagged and observed every three hours throughout anthesis. We initially assessed stigmatic peroxidase activity using Peroxtesmo KO paper (MacheryNagel, Düren, Germany). However, since peroxidase activity was detected as early as the bud stage, stigmatic exudate secretion was instead used to assess the pistillate phase onset, while the end of the pistillate phase was determined by stigma abscission. The onset of the staminate phase was delineated by anther dehiscence, with petal abscission marking the breakdown of the floral chamber and thus the end of the staminate phase. The methods for assessing thermogenesis are detailed in Method S1.

### Floral visitors and probable effective pollinators

Pollinator observation was conducted on 42 days across three years in Bowling Green Bay National Park (*ca.* 9h a day; 9^th^–15^th^ Dec, 2013) and Dingo Beach (*ca.* 6h a day; 7^th^–20^th^ Dec, 2018, 17^th^ January–6^th^ February, 2020), with a cumulative observation period of *ca.* 270 hours, with one observer per season. Because *Meiogyne* flowers have a tightly enclosed floral chamber, exhaustive inspection of flowers in the populations (by pulling apart the petals), as well as long-duration observation on designated plants were performed. Hourly observations (each lasting *ca.* 30 min) were conducted for the first 24 hours. Due to the lack of floral visitors and human-perceivable floral scent at night, subsequent observations were restricted to daytime. Floral visitors were preserved in either 50% isopropanol for identification, or frozen in -20°C and then silica-gel dried for assessing the presence of pollen. Probable effective pollinators were determined according to the following criteria: (1) visits to both sexual phases; (2) access to the inner floral chamber; and (3) presence of pollen grains on pistillate-phase floral visitors. Pollen grains on the floral visitors and the anther were examined using a Hitachi S3400 variable pressure scanning electron microscope (SEM). *Meiogyne heteropetala* has monosulcate monad pollen, which differ morphologically from other co-occurring flowering species (see also Method S1).

The activities of *Loberus sharpi* (the most likely effective pollinator) outside the flowers were observed in the vicinity to assess their interactions with potential models, revealing that *L. sharpi* is an aerial litter specialist. The petals were examined for the presence of eggs and larvae to assess whether the pollinators oviposit on the flowers.

### Characterisation of dimension and spectral reflectance

The dimensions of flowers (Fig. 1) and aerial litter (Fig. 1b, S1a–e) in the Dingo Beach and Conway National Park populations were measured using a digital caliper with the same approach: the first axis (length) was measured as the distance of the longest axis, with the subsequent axes (width & thickness) representing the next longest perpendicular to the previous axes. Because the outer corolla whorl encases almost all of the remaining floral organs and thus represents the major visual unit of the flowers, the spectral reflectance (300–700 nm) of the abaxial surface of outer petals and aerial litter were measured using an Ocean Optics FLAME-S-UV-VIS- ES spectrometer (Dunedin, FL, USA), a DH-mini UV-VIS-NIR lightsource and an R200-7-UV-VIS probe positioned at 45°. Individual flowers and aerial litter were used as replicates.

**Fig. 1.**
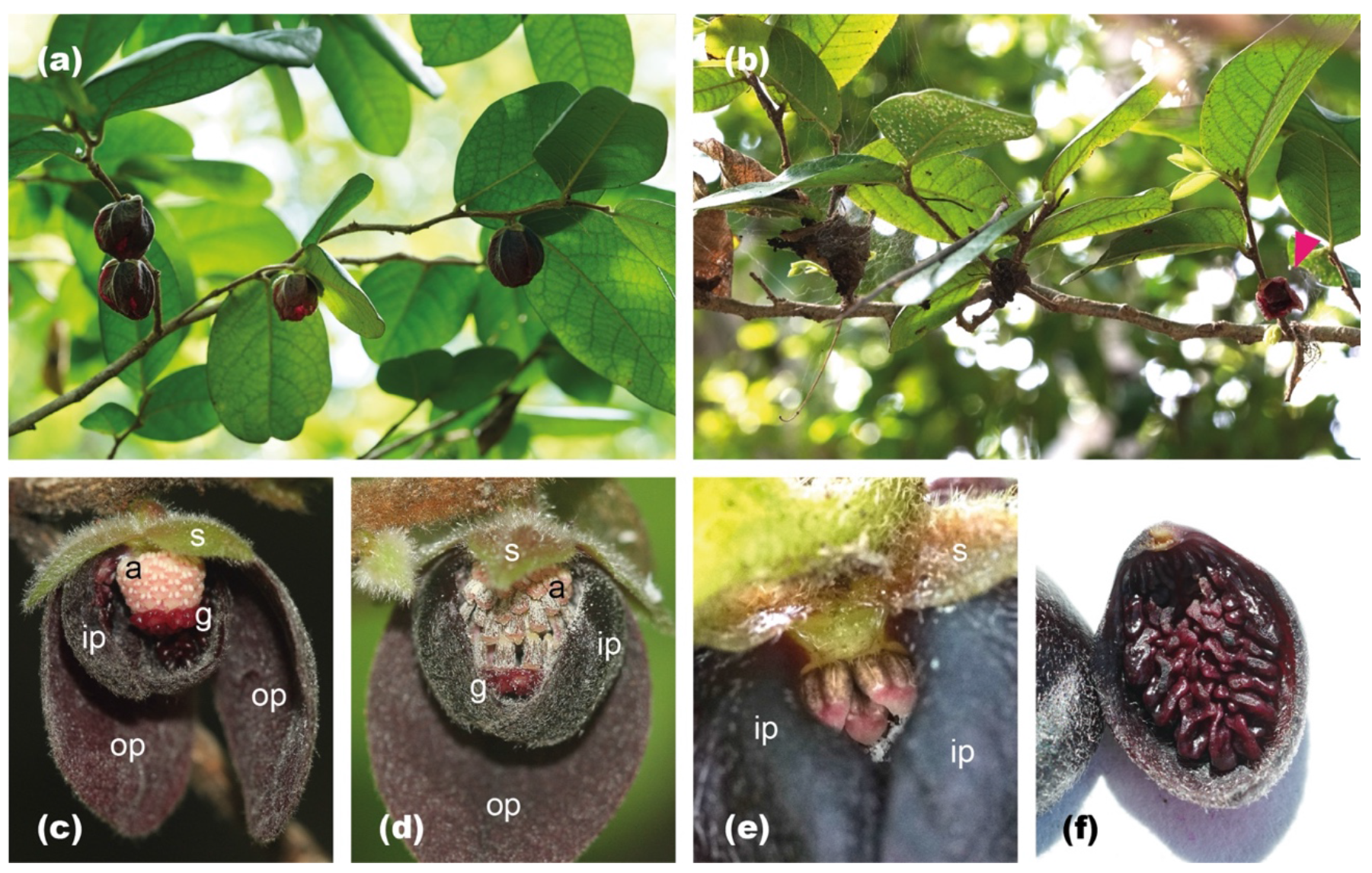
Flowers of *Meiogyne heteropetala*. (a) Axillary, solitary, and pendent flowers. (b) Aerial litter and *M. heteropetala* flower (latter indicated by a magenta arrowhead). (c) Pistillate-phase flower (one inner petal and one outer petal removed). (d) Staminate-phase flower (one inner petal and two outer petals removed). Stigmas and stamens abscised. (e) Basal aperture created by the gap between two adjacent inner petals (outer petals removed). (f) Inner petal with verrucose growth. a: androecium, g: gynoecium, ip: inner petal, op: outer petal, s: sepal.

### Floral odour characterisation

Due to the limited number of solid-phase microextraction (SPME) fibres, floral scent collection was restricted to the Dingo Beach population collected in 2018 and 2020. For the floral samples, each biological replicate consisted of five flowers from one individual; for the aerial litter samples, each biological replicate consisted of 10–30 pieces (*ca.* 10–20 g) of aerial litter shed from the same tree. Aerial litter was collected with the aim to sample as many common co-occurring tree species as possible. In total, biological replicates of pistillate-phase flowers (*n*=7), staminate-phase flowers (*n*=6), other *M. heteropetela* organs viz. flower buds, fruits, stem, and leaves (*n*=1–9), and aerial leaf (*n*=44) and twig (*n*=9) litter were collected. The buds, fruits, stem and leaves of *M. heteropetela* were mechanically damaged to maximise volatile release (otherwise too weak for detection). To gauge the general floral odours of the genus, we collected floral scents of three closely related Australian *Meiogyne* species with the typical floral syndrome of the genus and the family (yellow corolla and fruity odour), viz. *Meiogyne cylindrocarpa*, *M. trichocarpa* and *M. verrucosa* in Garry and Nada Sankowsky’s arboretum (in 2020, 2023; Tolga, Australia) and Gardens by the Bay (in 2018; Singapore) with the same sampling strategies. The organs were placed within a “Toppits” headspace bag. The headspaces were allowed to equilibrate for 30 min, followed by 30 min of volatile adsorption using a SPME fibre coated with polydimethylsiloxane-divinylbenzene (PDMS-DVB; 65-μm film thickness; Supelco, USA). To reduce contamination, the SPME fibres were sealed in double-layered headspace bags during transportation to the laboratory at the University of Hong Kong. Air in empty bags was also sampled at the field sites for both years to identify contaminants in ambient and transport environments. All compounds found in both the air control and the samples were omitted. Analyses were completed within three to five days post-extraction.

Dynamic headspace extraction was also performed to assess floral scent emission rate and composition variation across the three populations. Intact flowers were placed in headspace bags with two slits, one end receiving charcoal-filtered fresh air (ORBO 32, Supelco), and the other connected to a Porapak-Q cartridge (DVB; ORBO 1103, 50/80, Supelco). Floral odour was collected using a Spectrex PAS- 500 vacuum pump at a flow rate of 100 m/min over 4 h. Similarly, biological replicates were collected (pistillate: *n*=12; staminate: *n*=12), and the air in empty bag was likewise sampled to identify background contaminants for each location which were then omitted. Compounds on cartridges were eluted with 1 ml HPLC-grade hexane (Sigma-Aldrich, USA).

Gas chromatography-mass spectrometry (GC-MS) was performed in an Agilent 6890N/5973 gas chromatograph-mass selective detector. Either the volatiles were thermally desorbed from the SPME coating at 230°C for 1 min, or 1 μl eluent was injected into the inlet at 230°C for 1 min, before injecting into a Cyclodex-B chiral column (30 m × 0.25 mm, 0.25 μm thick film; J&W, USA) in splitless mode. Helium was used as the carrier gas at a flow rate of 1 ml/min. Oven temperature was ramped up from 40°C to 120°C at 2°C/min, followed by a ramp to 200°C at 50°C/min, which was held for 2 min. Analyses using a second DB-WAX column (20 m × 0.255 mm, 0.25 μm thick film; J&W) for both SPME and dynamic headspace samples were performed with identical inlet and carrier gas settings. Oven temperatures were held at 60°C for 3 min, followed by a ramp up to 250°C at 10°C/min, and finally held for 7 min. Peaks were manually integrated and tentatively identified using the Wiley and National Institute of Standards and Technology mass spectral libraries (NIST, USA; 2014), with a threshold of 85% library search best fits. Compound identity was assessed against published standardised retention index values or verified by co-injecting with commercially available standards. Relative abundances were calculated by measuring area under curve of the integrated peaks. Odour similarity of the SPME data was visualised using two-dimensional non-metric multidimensional scaling (NMDS) in the R package vegan (Oksanen et al., 2013). Unidentified compounds were excluded. When stereochemistry was unclear, isomers were integrated. Bray-Curtis distances were calculated using square-root transformed data. ANOSIM statistics were performed with 10,000 random permutations to assess differences in scent composition between predefined groups. Post-hoc SIMPER was used to identify volatiles responsible for difference between groups, with 999 permutations and a cut-off point at 1% contribution.

### RNA extraction

Due to logistical limitations, the plant tissues were preserved in RNAlater (Invitrogen, USA), and promptly stored and transported at -20°C (biological replicates per organ from 4–9 individuals; each replicate consists of one individual; collected at 12–2pm in Dingo Beach, 2020). To ensure full permeation of RNAlater, plant tissues were finely diced immediately before fixation. Total RNA was extracted using the CTAB method. Tissues were ground and incubated in CTAB buffer (2% CTAB, 2% PVP, 100 mM Tris-HCl, 2 M NaCl, 2% β-mercaptoethanol, pH 8.0) for 25 min at 65°C. Cell debris was removed by centrifugation and the supernatant washed with chloroform:isoamyl alcohol (24:1) twice, before precipitation in 2.5 M LiCl overnight at 4°C. The RNA pellets were resuspended in SSTE buffer (1 M NaCl, 0.5% SDS, 10 mM Tris-HCl, 1 mM EDTA, pH 8.0), washed once with chloroform:isoamyl alcohol, and precipitated in 70% ethanol. After rinsing in 70% ethanol, the air-dried RNA pellets were eluted in H2O.

### Transcriptome

The procedure for generating the floral transcriptome is detailed in Methods S1. Total mRNA of a pistillate phase flower was captured using oligo(dT) method before fragmentation. cDNA was synthesised and tagged with adaptors, followed by size selection by agarose gel electrophoresis and PCR enrichment. After library preparation, RNA-seq was performed on Illumina Hi-seq 4000 platform at Beijing Genomics Institute. Contigs were assembled using Trinity v2.0.6 (Grabherr et al., 2011) and annotated with NT, NR, GO, KOG, KEGG, SwissProt and InterPro databases. Clean reads were mapped onto contigs using Bowtie2 v2.2.5 (Langmeads & Salzburg, 2012). Fragments per kilobase of transcript per million reads (FPKM) were calculated using Cufflinks (Trapnell et al., 2010). KAAS (Moriya et al., 2007) was used to assign KO identifiers and map the contigs to KEGG pathways, allowing identification of contigs involved in anthocyanin and terpene production. The raw data was deposited at NCBI SRA database (accession number: PRJNA972425).

### Reverse-transcription quantitative PCR (RT-qPCR)

Among the TPS contigs, five contigs with full length coding sequence (CDS) were identified. The relative expression level of these contigs with full length CDS was assessed by RT-qPCR. Primers were designed such that: (1) annealing temperature is *ca.* 60°C; (2) amplicons are *ca.* 150–200 bp long; and (3) at least one primer of each pair spans across an exon-intron boundary at the 3’ end (Table S1). To synthesise cDNA, 2 µg of total RNA were reverse transcribed using High-Capacity cDNA Reverse Transcription Kit with RNase Inhibitor (Applied Biosystems, USA). Quantitative PCR was then performed using PowerUp SYBR Green Master Mix (Applied Biosystems) under Bio-rad CFX96 system, which automatically generates Ct values. The reaction mixtures were kept at 50°C for 2 min, and then heated to 95°C for 2 min, followed by 40 cycles of 15 s denaturation at 95°C and 1 min annealing/extension at 60°C. Melting curve analyses were performed to confirm single amplicon formation. Relative expression levels were calculated using the Pfaffl method (Pfaffl, 2001). The geometric means of two housekeeping genes ACT2 and UBC21 were used for normalisation (Vandesompele et al., 2002). The non-parametric Kruskal-Wallis test and post-hoc Dunn test were performed, with Benjamini-Hochberg correction applied for multiple comparisons.

### Bacterial expression of terpene synthase

The CDS of TPSs were amplified by Phusion polymerase (New England Laboratory, USA) from floral cDNA. Amplification was successful for only one TPS. The CDS was then inserted into pET29 plasmid, with the addition of 6×His tag at the C-terminal. The sequence upstream of the RRX8W motif, including the transit peptide, was deleted for activation of monoterpene synthase (Williams et al., 1998). The vector was then transformed into Rosetta cells (Novagen, Germany). The clone was cultured at 37°C with 100 µg/ml ampicillin and 25 µg/ml Chloramphenicol until OD600 reached 0.5–0.7. TPS expression was induced at 4 µM IPTG (Invitrogen) for 20 hours at 21°C. Cells were harvested and lysed in 1 mg/ml lysozyme buffer (50 mM NaH2PO4, 300 mM NaCl, 10mM imidazole, pH 8.0) on ice for 30 min, followed by sonication. Lysates were mixed with Ni-NTA slurry (Qiagen) in 1:4 ratio and incubated at 4°C for 60 min, before being washed twice in ice-cold wash buffer (50 mM NaH2PO4, 300 mM NaCl, 20mM imidazole, pH 8.0). The purified protein was eluted in buffer (50 mM NaH2PO4, 300 mM NaCl, 250mM imidazole, pH 8.0). Protein concentration was measured using BCA test (Pierce, ThermoFisher, USA). The purified enzyme was mixed with equal volume of glycerol and stored at -20°C. All downstream experiments were performed within three days. The primers used are listed in Table S1.

### Functional characterisation of TPS

MnCl2 and MgCl2 concentration and pH for the enzyme reaction were initially optimised. To determine enzyme kinetic parameters and characterise product profile, 0.15 ml enzyme extract was mixed with 1.35 ml assay buffer (50 mM MOPSO, 50 mM MgCl2, 5 mM DTT, 10% (v/v) glycerol, pH 7.0) in a 2 ml glass vial. Reaction mixtures were overlaid with 100 µl hexane for terpenoid extraction. Enzymatic reaction was initiated by the addition of 1.25–80 µM (enzyme kinetics) or 500 µM (product profile characterisation) geranyl pyrophosphate (GPP; Echelon Biosciences, USA) with an incubation period of 2 hours at 25°C. For assaying sesquiterpene synthase activity, 54 μM farnesyl pyrophosphate (FPP; Echelon Biosciences) was added instead. The hexane layer was removed for GC-MS analyses. All treatments were triplicated.

### Y-tube olfactometer bioassay

To study the behavioural responses of beetle pollinators to odour sources/blends, a two-choice Y-tube olfactometer (WT-1018; Sigma Scientific) was used. The experiment was conducted in a dim room at 25°C. An air-pump was connected to the base of the olfactometer to create a flow at 100 ml/min. The two streams of incoming air were humidified and charcoal-filtered (ORBO 32; Supelco). For plant material, one flower or one piece of aerial leaf litter was placed in an arm. For synthetic mix experiments, 300 ng of total synthetic chemicals were mixed, based on the ratio of volatiles emitted by the flowers, or the products of the enzymatic reaction (Table S2). After the beetles were collected from aerial litter in the wild, they were starved for 1 h. The assay was initiated by the release of the beetle into the olfactometer. A choice was made when the beetle entered either arm 30 mm from the split and stayed for at least 1 min. A ‘no choice’ was recorded when it did not decide within 5 min. The arms were interchanged every five tests to remove positional bias. The flowers and aerial litter were replaced with a fresh set after every 10 tests. A total of 68 beetles were used in bioassays. While the beetles were not reused in the same bioassay to ensure independent sampling, they were subjected to multiple different bioassays owing to limited availability (32 beetles for three assays for plant materials; another 36 beetles for five assays for synthetic mixes). Generalised linear mixed models (GLMM) were performed to account for re-use of beetles (see Method S1). A binomial test was also conducted to detect odour preferences.

## Results

### Floral architecture and phenology

*Meiogyne heteropetala* flowers have two whorls of three dark maroon petals that superficially resemble aerial leaf litter (Fig. 1a, b). The papery outer petals form a loose outer floral chamber. The strongly concave and fleshy inner petals are imbricate and tightly pressed against each other, restricting the access of the inner chamber to three basal apertures (2.6 ± 0.5 × 2.4 ± 0.4 mm, *n* = 24; Fig. 1e) and effectively creating a size filter for pollinators. The inner petals possess an elaborate verrucose basal growth, emanating from a pocket-like tissue growing along longitudinal ridges that extends over almost the entire adaxial surface (Fig. 1f). The flowers are bisexual and protogynous (Fig. 3a). Anthesis extends over *ca.* 5 days, with a 2-day pistillate phase and a 3-day staminate phase that overlap by 1–3 hours. Scent emission detectable by human perception is restricted to daytime and peaks *ca.* 12–2pm. The flower remains pendent with minimal petal movement throughout anthesis. No thermogenesis was detected. Due to time and cost constraint, autogamy and apomixis were not assessed, but the Annonaceae are predominantly outcrossing, with no reports of obligate self-fertilisation (Pang & Saunders, 2014; Saunders, 2020).

### Floral visitors and probable effective pollinators

Arthropod visitation to *M. heteropetala* flowers was infrequent. In Dingo Beach, a total of 193 floral visitors were recorded in 2018 and 2020, consisting of the erotylid beetle *Loberus sharpi* (Fig. 2; 8%; *n*=16), cockroach nymphs (49%; *n*=94), spiders (37%; *n=*72), Pseudococcidae mealybugs (2%; *n*=3), *Crematogaster* ants (2%; *n*=4), ricanid planthoppers *Scolypopa australis* (1%; *n*=2) and a katydid nymph (1%; *n*=2). Only two chloropidae flies were observed visiting the flowers in the Bowling Green Bay National Park in 2013. We were unable to identify the cockroach visitor due to difficulty in nymph identification. All visitors except the planthoppers and katydid were recorded visiting both sexual phases. The inner floral chamber, which encloses the reproductive organs, was only accessible to *L. sharpi* and early instar cockroach nymphs. Under SEM, no pollen was detected on the fifteen cockroach nymphs collected from the inner floral chamber of pistillate- phase flowers. However, pollen grains were observed on four out of seven *L. sharpi* individuals collected from the inner floral chamber of pistillate-phase flowers, suggesting that it is the most likely effective pollinator (0–38 pollen grains per individual; Fig. 2g–j). Visitation by *L. sharpi* was very low in *M. heteropetala*. Only 16 individuals were observed over 270 h across three years. Visitation rate also appeared to be highly sensitive to temporal and geographical variation (unusually low in Bowling Green Bay National Park in 2013). The fruit set was noticeably low towards the end of the flowering seasons, suggesting that apomixis is unlikely, and is consistent with the low visitation rate of *L. sharpi*. It is unclear whether *L. sharpi* consumes pollen or petal corrugation. No bite marks, however, were observed on the inner petal corrugations.

**Fig. 2.**
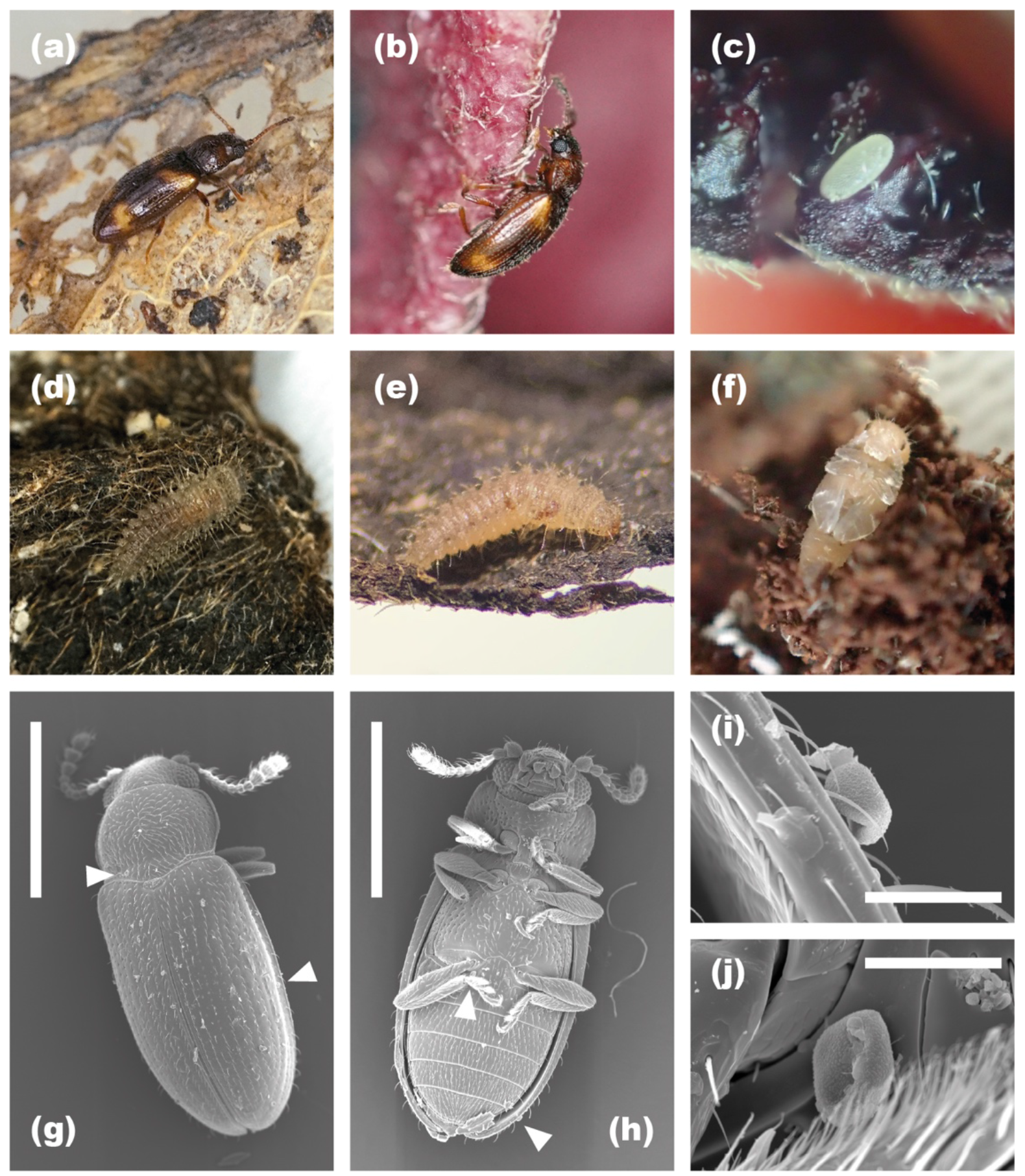
Probable effective pollinator *Loberus sharpi.* (a) Adult in aerial leaf litter. (b) Adult on an outer petal of *Meiogyne heteropetala*. (c) Egg laid in the crevice of the inner petal corrugation of *M. heteropetala*. (d, e) Larvae reared in laboratory on dried *M. heteropetala* petals (d: early instar; e: late instar). (f) Pupa reared in laboratory on decayed inner petal mixed with frass. (g–j) Scanning electron micrographs of *L. sharpi*. (g) Dorsal view. (h) Ventral view. (i) Pollen grain on the left elytron. (j) Pollen grain deposited on a sternite. Scale bars: (g), (h) 0.5 mm; (i), (j) 50 µm.

**Fig. 3.**
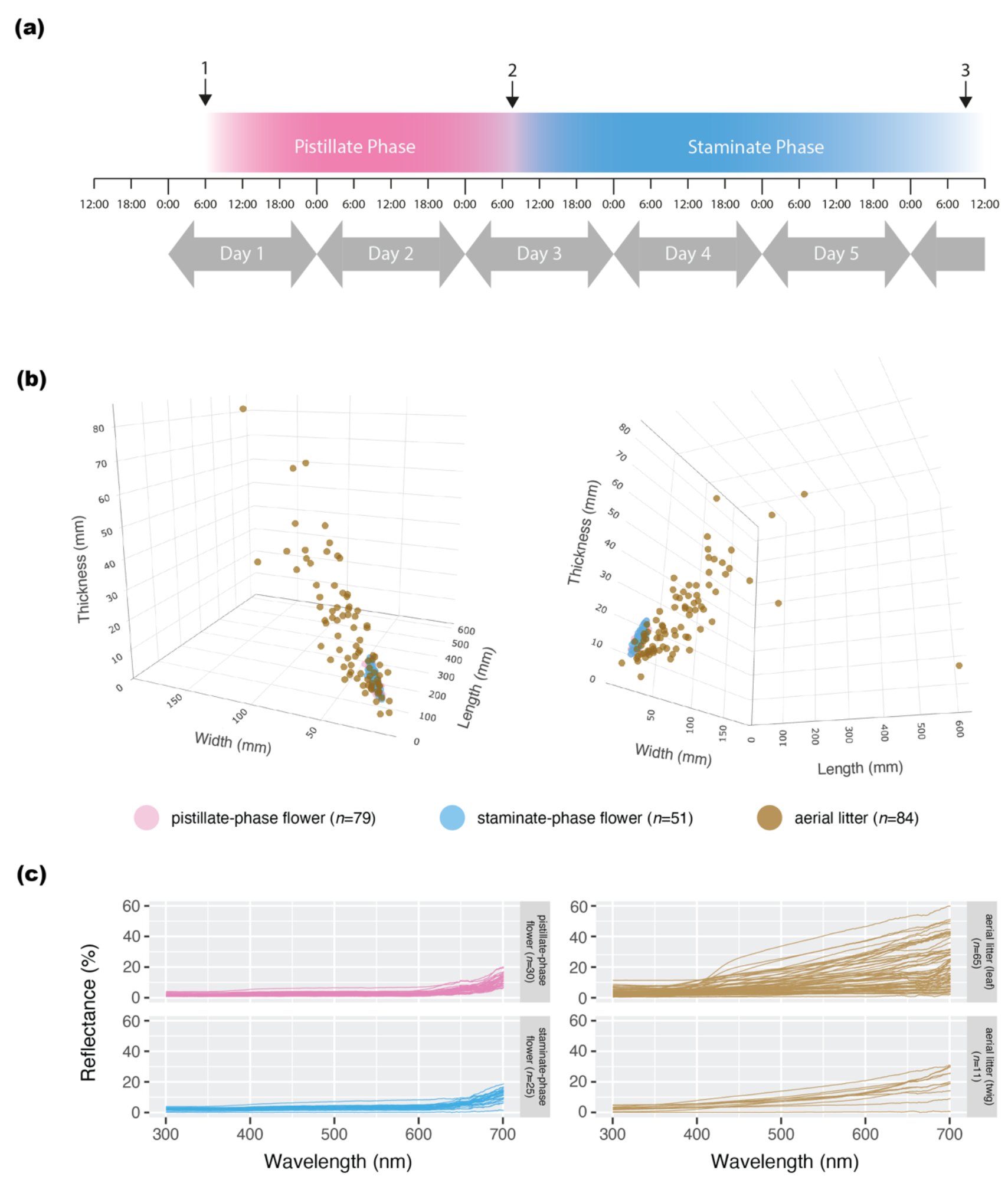
Characterisation of floral ontogeny and visual cues of *Meiogyne heteropetala* flowers and aerial litter. (a) Floral phenology. 1: Onset of pistillate phase. 2: anther dehiscence, stigma abscission and stamen abscission (with the three events occurring in chronological order within 1–3 hours). 3: earliest observed petal abscission. (b) 3D plots of dimensions of pistillate-phase flowers (*n*=79), staminate- phase flowers (*n*=51) and aerial litter (*n*=84) from two angles. (c) Spectral reflectance of pistillate-phase flowers (*n*=30), staminate-phase flowers (*n*=25), aerial leaf litter (*n*=65) and aerial twig litter (*n*=11).

*Loberus sharpi* is likely an aerial litter specialist. Adults (*n=*115) and larvae (*n=*16) were found in the abundant co-occurring aerial leaf litter on multiple tree and undergrowth species, sometimes intermixed with dead twigs. The beetles were absent on co-occurring flowers, fruits, branches, bark, or ground leaf litter.

A total of 15 eggs were observed in the crevices of the inner petal corrugation during anthesis (Fig. 2c). The vast majority of petals fell onto the ground upon abscission. No larvae were observed from 50 fallen petals inspected, suggesting that the larvae likely could not survive in fallen petals on the forest floor. In the unlikely event that the abscised corolla remained arboreal, larvae of all stages were observed on the aerial dried petals. In the laboratory, the eggs in inner petals hatched and pupate to adults relying solely on the abscised petals (Fig. 2d–f). The morphology of pupated adults and larvae were congruent with *L. sharpi* adults collected in the field and published *Loberus* larvae (Carlton et al., 2000) respectively.

In the two locations assessed, 20–20.6% of aerial litter clusters were free-hanging on branches (Fig. S1), while the remaining 79.4–80% were anchored by some agents, including 18–23.5% self-retained by vasculature, 44.1–48% anchored by arthropod silk, 0–8.8% by fungal hyphae, 2–2.9% by tendrils, 0–12% by mixed agents (Fig. S1). Common arthropods that secure aerial litter with silks include green tree ants, spiders, and moth larvae, all of which were observed to utilise aerial litter as brood shelters.

### Dimension and spectral reflectance

Aerial litter clumps vary greatly in size (Fig. 3b), while *Meiogyne heteropetala* flowers vary much less in size. PERMANOVA test detected a significant size difference among pistillate-phase flowers, staminate-phase flowers, and aerial litter clumps (dF: 2; SST: 6.649; *R*^2^: 0.512; *F-*value: 110.72; *p*-value: 0.001***). A significant difference was detected between the flowers (pistillate+staminate) and aerial litter clumps (dF: 1; SST: 6.641; *R*^2^: 0.512; *F-*value: 222; *p*-value: 0.001***), but there was no difference between the sexual phases (dF: 1; SST: 0.00707; *R*^2^: 0.01116; *F-*value: 1.4446; *p*-value: 0.223, ns), nor populations (dF: 1; SST: 0.00725; *R*^2^: 0.01146; *F-*value: 1.4838; *p*-value: 0.211, ns). However, the 3D plots (Fig. 3b) and PCA plot (Fig. S2) showed the flowers occupied a space considerably overlapping with the smaller end of aerial litter space.

Likewise, the spectral reflectance of aerial litter was more variable than the flowers (Fig. 3c). The spectral reflectance of the flowers was similar across populations and sexual phases, and matched with a subset of aerial litter (Fig. 3c), suggesting advergence of visual cues. Unfortunately, existing data on erotylid vision systems in the literature are insufficient for reconstructing colour loci.

### Flower and aerial litter scent composition

We identified 29 floral volatiles (Table 1) belonging to monoterpenes or sesquiterpenes, with one molecule unknown. The most abundant class was monoterpenes and monoterpenoids (pistillate: 96.72 ± 12.7%; staminate: 97.34 ± 0.63%). 1,8-cineole was the most abundant monoterpene (pistillate: 76.28 ± 1.6%; staminate: 79.22 ± 0.93%), followed by (+)-α-pinene (pistillate: 4.01 ± 0.25%; staminate: 3.77 ± 0.31%) and (-)-limonene (pistillate: 3.62 ± 0.35%; staminate: 3.2 ± 0.19%). Four sex-specific compounds were found at low abundance (< 0.2%).

**Table 1.** Floral scent composition of *Meiogyne heteropetala* derived from SPME-GC-MS and MhCINS product profile measured by area under curve in TICs. Compounds in bold were verified by commercially available standards. Asterisks indicate compounds also found in aerial litter headspaces (see also Table S3). “…” denotes the absence of the molecule in the samples.

|  | Kovats index |  | relative abundance (%) |  | MhCINS<br>(n=4) |
| --- | --- | --- | --- | --- | --- |
|  | Cyclodex-B | DB-WAX | pistillate-phase<br>flower (n=7) | staminate-phase flower<br>(n=6) |  |
| **Monoterpenes & Monoterpenoids** |  |  |  |  |  |
| α-thujene* | 960 | 1016 | 0.56 ± 0.07 | 0.46 ± 0.06 | 0.39 ± 0.03 |
| **(-)-α-pinene\*** | 1001 | 1017 | 0.22 ± 0.02 | 0.20 ± 0.02 | 0.13 ± 0.04 |
| **(+)-α-pinene\*** | 1009 | 1017 | 4.01 ± 0.25 | 3.77 ± 0.31 | 5.62 ± 0.35 |
| **β-myrcene\*** | 1023 | 1155 | 2.07 ± 0.35 | 1.32 ± 0.36 | 1.10 ± 0.22 |
| **sabinene\*** | 1027 | 1111 | 2.77 ± 0.39 | 2.96 ± 0.17 | 2.16 ± 0.16 |
| β-thujene | 1036 | 1208 | 0.16 ± 0.03 | 0.17 ± 0.02 | 0.03 ± 0.03 |
| **(-)-α-phellandrene\*** | 1049 | 1156 | 0.10 ± 0.02 | 0.08 ± 0.01 | ... |
| α-terpinene* | 1051 | 1174 | 0.35 ± 0.08 | 0.27 ± 0.06 | ... |
| (+)-β-pinene* | 1053 | 1105 | 1.35 ± 0.11 | 1.35 ± 0.08 | 1.17 ± 0.07 |
| **(-)-β-pinene\*** | 1061 | 1105 | 0.26 ± 0.02 | 0.23 ± 0.02 | 0.11 ± 0.05 |
| **cis-β-Ocimene\*** | 1071 | 1239 | 0.12 ± 0.02 | 0.09 ± 0.02 | 0.02 ± 0.02 |
| **(-)-limonene\*** | 1073 | 1196 | 3.62 ± 0.35 | 3.20 ± 0.19 | 0.84 ± 0.11 |
| **(+)-limonene\*** | 1077 | 1196 | 2.87 ± 0.33 | 2.29 ± 0.19 | 0.91 ± 0.05 |
| **trans-β-ocimene\*** | 1083 | 1256 | ... | ... | 0.07 ± 0.04 |
| (+)-β-phellandrene* | 1088 | 1214 | 0.22 ± 0.12 | 0.04 ± 0.04 | 0.10 ± 0.04 |
| (-)-β-phellandrene* | 1093 | 1214 | 0.37 ± 0.07 | 0.30 ± 0.03 | 0.18 ± 0.06 |
| **γ-terpinene\*** | 1102 | 1249 | 0.76 ± 0.14 | 0.50 ± 0.13 | 0.12 ± 0.04 |
| **terpinolene\*** | 1123 | 1283 | 0.22 ± 0.03 | 0.16 ± 0.04 | 0.02 ± 0.02 |
| **1,8-cineole\*** | 1128 | 1210 | 76.28 ± 1.60 | 79.22 ± 0.93 | 80.45 ± 0.54 |
| (-)-linalool* | 1253 | 1554 | ... | ... | 0.80 ± 0.18 |
| (+)-linalool* | 1256 | 1554 | ... | ... | 1.07 ± 0.21 |
| **(-)-α-terpineol\*** | 1378 | 1711 | 0.12 ± 0.05 | 0.33 ± 0.18 | 2.17 ± 0.25 |
| **(+)-α-terpineol \*** | 1379 | 1711 | 0.27 ± 0.08 | 0.58 ± 0.25 | 2.52 ± 0.30 |
| **Sesquiterpenes & Sesquiterpenoids** |  |  |  |  |  |
| β-calarene | 1407 | 1602 | ... | 0.19 ± 0.07 | ... |
| **(E)-β-caryophyllene\*** | 1489 | 1608 | 1.22 ± 0.57 | 0.76 ± 0.21 | ... |
| aromandendrene* | 1498 | 1620 | 0.38 ± 0.22 | 0.35 ± 0.12 | ... |
| germacrene D | 1500 | 1729 | 0.47 ± 0.35 | 0.21 ± 0.09 | ... |
| **humulene\*** | 1512 | 1681 | 0.03 ± 0.03 | ... | ... |
| γ-muurolene* | 1530 | 1700 | 0.16 ± 0.11 | ... | ... |
| Bicyclogermacrene* | 1552 | 1749 | $0.35 \pm 0.22$ | $0.41 \pm 0.14$ | ... |
| **Unknown** |  |  |  |  |  |
| 56(29), 41(23), 43(16),<br>42(12), 55(6), 39(6),<br>57(2), 40(2), 45(2),<br>44(1) | 838 | 980 | $0.67 \pm 0.20$ | $0.54 \pm 0.08$ | ... |
| **Monoterpenes & Monoterpenoids** | | | $96.72 \pm 12.7$ | $97.34 \pm 0.63$ | $100 \pm 0.00$ |
| **Sesquiterpenes & Sesquiterpenoids** | | | $2.61 \pm 1.11$ | $2.12 \pm 0.60$ | ... |
| **Unknown** | | | $0.67 \pm 0.18$ | $0.54 \pm 0.09$ | ... |

The headspace of aerial litter was highly heterogenous and consisted of 159 volatiles from six classes (Table S3; Fig. 5b), of which the largest two were sesquiterpenes and sesquiterpenoids (leaf: 53.79 ± 4.40%; twig: 60.47 ± 10.74%) and monoterpenes and monoterpenoids (leaf: 43.79 ± 4.73%; twig: 36.37 ± 11.4%).

There was no volatile ubiquitously detected in all aerial litter samples. Although the aerial litter on average had a lower 1,8-cineole content (leaf: 1.70 ± 0.55%, ranges 0–17.5%; twig 3.99 ± 2.93%, range: 0–27.04%), it shared a similar monoterpenoid upper range with flowers (aerial litter: 98.59%; flowers: 99.17%). 1,8-cineole was ranked top 15.7% most common volatile (by frequency), and top 8.2% most abundant volatile (by relative abundance) in aerial litter (Table S3). The flowers, moreover, shared 25 out of 29 (86%) volatiles with aerial litter, constituting 98.7% (pistillate-phase) and 98.9% (staminate-phase) of floral VOCs in relative abundance (Table 1).

Scent emission rate of pistillate- and staminate-phase flowers was 83.7 ± 22.5 ng/min and 147 ± 35.15 ng/min respectively. Overall floral scent emission rate ranged from 2.1–418.9 ng/min. Aerial leaf litter had an emission rate of 78 ± 10.2 ng/min, while aerial twig litter had an emission rate of 50.3 ± 17.5 ng/min. Scent emission rate of aerial litter ranged from 23.9–91.9 ng/min. There was no significant difference in emission rate among flowers and aerial litter (Kruskal-Wallis test, χ^2^ = 1.50, df = 3, *p*=0.68). Multiple linear regression suggested there was no difference in emission rate among populations (*p*=0.515), nor between sexes (*p*=0.383). There was no difference in scent profile across populations, with small R value (Fig. S3; ANOSIM R=0.093; *p*=0.066).

Aerial leaf litter odour composition was strongly dependent on the species the leaf was shed from (ANOSIM R: 0.996; *p*<0.001***). The odour of fresh *M. heteropetala* tissues and the aerial litter derived from *M. heteropetala* are highly similar: 83.12 ± 1.68% of *M. heteropetala* aerial leaf litter volatiles in relative abundance were also detected in fresh *M. heteropetala* leaves (Table. S3). Similarly, 87.33% of *M. heteropetala* aerial twig litter volatiles were also detected in fresh *M. heteropetala* stems (Table. S3). Therefore, aerial litter VOCs were likely the result of slow release of endogenously produced terpenoids built up before senescence. We, however, cannot rule out the microbial influence on litter odour: we suspect, at the very least, microbial degradation would weaken litter structural integrity and may facilitate volatile release.

Aliphatic ester was the most abundant class of floral volatiles for the three congener (Table S3), *M. cylindrocarpa* (98.32 ± 0.42%), *M. trichocarpa* (95.23 ± 1.98%) and *M. verrucosa* (99.12%), including isoamyl acetate (*M. trichocarpa*: 95.23 ± 1.98%; *M. cylindrocarpa*: 72.73 ± 7.67%), isobutyl acetate (*M. cylindrocarpa*: 24.03 ± 6.35%), and ethyl isovalerate (*M. verrucosa:* 99.12%). *Meiogyne cylindrocarpa* and *M. trichocarpa* were visited by nitidulid and curculionid beetles (unpublished data). The yellow corolla and fruity odour aligned with the typical Annonaceae floral syndrome (Goodrich, 2006) and likely represent the symplesiomorphic states in *Meiogyne*.

### Visualisation of olfactory space

Because monoterpenes are common plant secondary metabolites found in both floral and vegetative organs, floral monoterpenes could be universally emitted compounds. To address the floral specificity of volatiles and potential sexual partitioning in floral odour space, organ headspaces were compared using NMDS (Fig. 4a; 2-D stress value: 0.054; ANOSIM global, all organs; R = 0.714; *p* = 0.0001***). The floral headspaces of both sexual phases overlapped and shared highly similar odour space, suggesting that there was no sexual differentiation in scent profile. Floral headspace furthermore occupied a distinct odour space from other non-floral tissues, indicating a clear floral specialisation in olfactory cues.

**Fig. 4.**
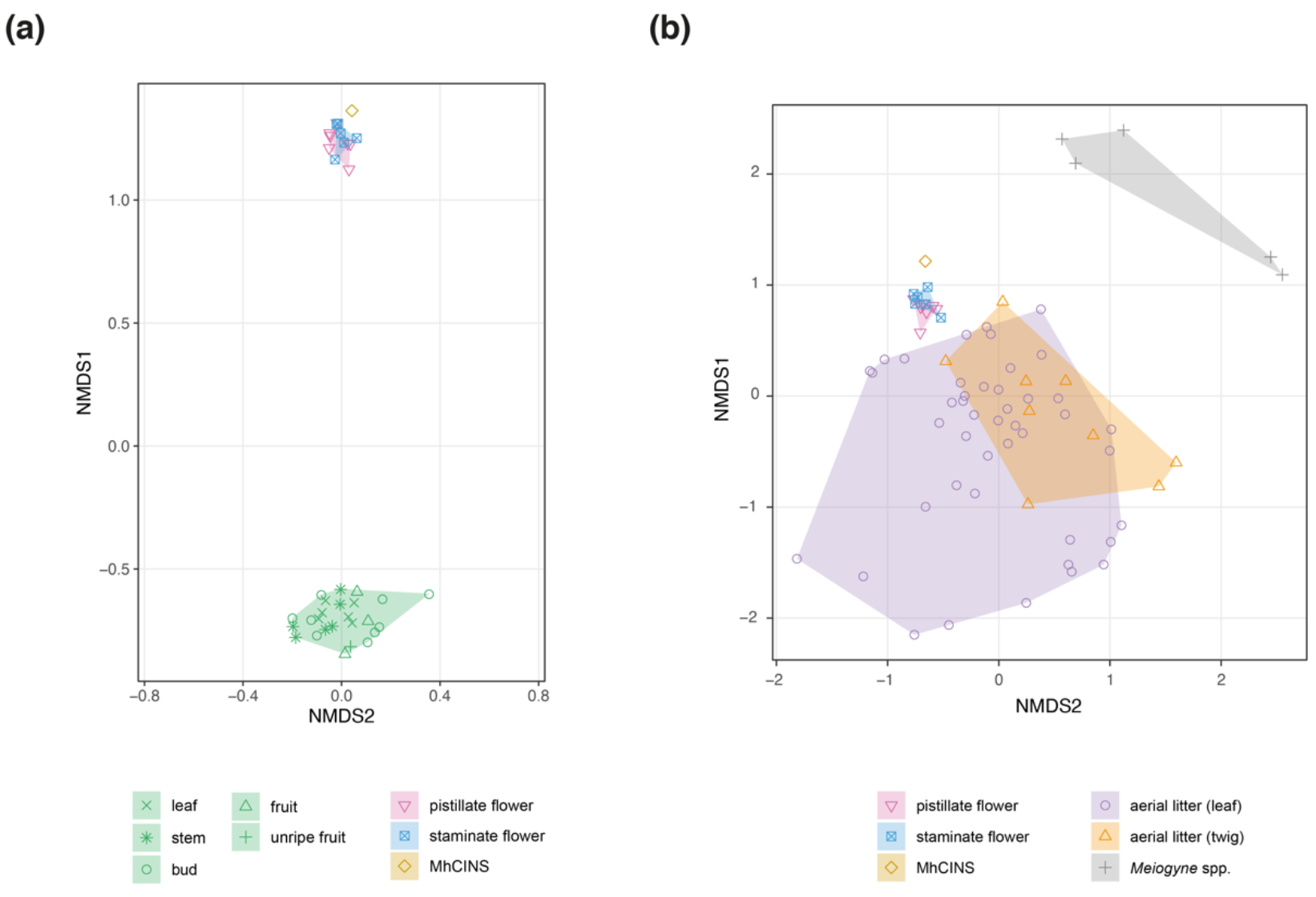
Visualisation of odour space. (a) Non-metric dimensional scaling (NMDS) based on Bray-Curtis dissimilarity index values of 156 volatiles of MhCINS and 38 samples of *Meiogyne heteropetala* organs, including pistillate-phase flowers (*n*=7), staminate-phase flowers (*n*=6), leaves (*n*=6), stems (*n*=6), flower buds (*n*=9), unripe fruit (*n*=1) and ripe fruits (*n*=3). 2-D stress value: 0.0543; ANOSIM global R=0.714; *p*=0.0001***. (b) NMDS of 77 odour sources, including MhCINS products, 13 samples of *M. heteropetala* flowers (pistillate-phase flowers (*n*=7); staminate-phase flowers (*n*=6), aerial leaf litter (*n*=44), aerial twig litter (*n*=9), and flowers of other *Meiogyne* species (*M. cylindrocarpa* and *M. trichocarpa*; *n*=5), based on Bray-Curtis dissimilarity index values of 157 volatiles. 2-D stress value: 0.14; ANOSIM global R=0.397; *p*=0.0002***.

**Fig. 5.**
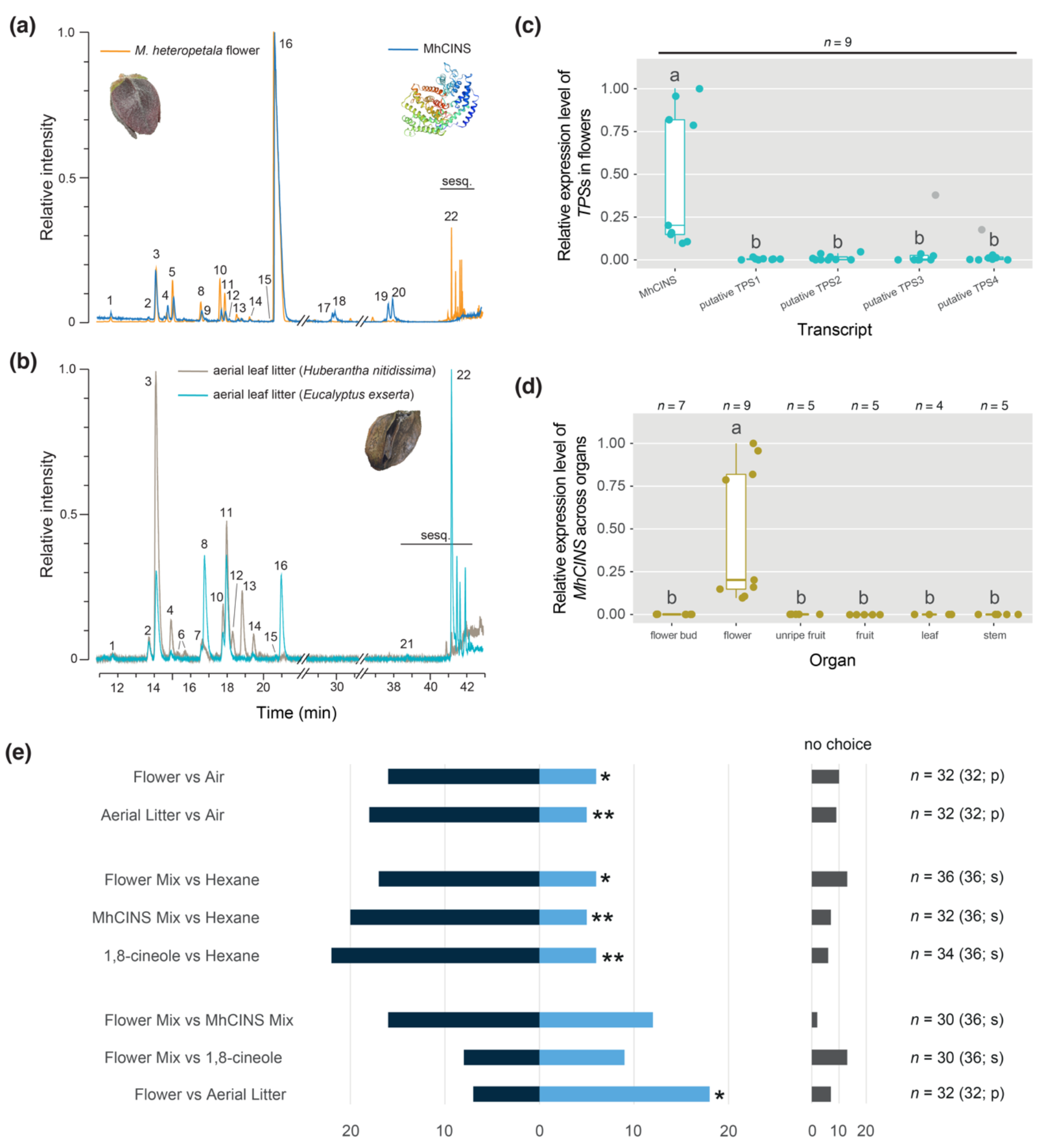
(a) Total ion chromatographs (TIC) of *Meiogyne heteropetala* floral scent (SPME) and MhCINS products with Cyclodex-B column. Homology model of MhCINS was reconstructed using SWISS-MODEL (see Method S1). (b) TIC of representative aerial litter scent (SPME) derived from the dead leaves *Huberantha nitidissima* and *Eucalyptus exserta* with Cyclodex-B column. (1: α-thujene; 2: (-)- α-pinene; 3: (+)-α-pinene; 4: β-myrcene; 5: sabinene; 6: camphene isomers; 7: (-)-α-phellandrene; 8: (+)-β-pinene; 9: (-)-β-pinene; 10: (-)-limonene; 11: (+)-limonene; 12: trans-β-ocimene; 13: β-phellandrene isomers; 14: γ-terpinene; 15:terpinolene; 16: 1,8-cineole; 17: (-)-linalool; 18: (+)-linalool; 19: (-)-α-terpineol;20: (+)-α-terpineol; 21: β-Maaliene; 22: (E)-β-caryophyllene; sesq.: sesquiterpenoids). (c) Relative expression level of terpene synthases in the flower by RT-qPCR (non-parametric Kruskal-Wallis test with Dunn post-hoc test). Different alphabets above the box and whisker denote significant difference after Benjamini-Hochberg correction. Grey dots denote outlier based on the 1.5 IQR rule. (accession no.: MhCINS: OQ592534; putative TPS 1–4: OQ805854–OQ805857). (d) Relative expression level of MhCINS across organs by RT-qPCR. (e) Two- choice bioassay responses from *L. sharpi* (binomial test). *n* refers the number of beetles subjected to the bioassay. Number in parentheses refers to the total number of beetles used in the pool. In total 68 beetles were used. p: plant material; s: synthetic mix. (*: *p*<0.05; **: *p*<0.01; ***: *p*<0.001).

To assess olfactory advergence to potential models, another NMDS was performed on the products of *Meiogyne heteropetala cineole synthase* (MhCINS; see next section), *M. heteropetala* flowers, aerial leaf litter and floral scent for other *Meiogyne* species (Fig. 4b; 2-D stress value: 0.14; ANOSIM global R = 0.397; *p* = 0.0002***). *Meiogyne verrucosa* constricted all other data points to a very tight cluster due to its disparity in odour composition, and was therefore excluded from the NMDS. The odour space defined by the leaf and twig litter is broad, representing a wide variation in odour blends. The floral scent samples clustered tightly, indicating little variation between samples. While the floral samples on the NMDS do not overlap the range of litter samples (Fig. 4b), the floral odour loci were very close to that of aerial litter. There are numerous litter samples that are more closely grouped with the floral scent than with other litter on the same plot. The close proximity of *M. heteropetala* floral scents to the aerial litter odours and much greater distance between the floral scents of *M. heteropetala* and other *Meiogyne* species provides evidence of olfactory advergence of *M. heteropetala* floral scent towards litter models.

In both NMDS plots, inclusion of unknown molecules yielded very similar olfactory landscapes. SIMPER (Table S4) reveals that six molecules were significantly responsible for the difference between *M. heteropetala* flowers and aerial litter, including 1,8-cineole (contribution %: 12.4%), (*E*)-β-caryophyllene (4.3%), sabinene (2.4%), aromandendrene (2.3%) and (-)-limonene (1.7%).

### Terpenoid and anthocyanin metabolism in the transcriptome

In the floral transcriptome, 31,269 contigs were detected (Fig. S4). Fifty-eight contigs from the anthocyanin biosynthetic pathway were identified (Fig. S5), among which four contigs were in the top 1% most highly expressed contigs in the transcriptome. For terpenoid biosynthesis, 18 contigs for MEP pathway and eight contigs for MEV pathway were retrieved in the floral transcriptome (Fig. S6). Among the six contigs of geranyl or farnesyl disphosphate synthases identified, one contig was among the top 1% most highly expressed contigs in the transcriptome.

Fifteen putative TPS contigs were identified, among which five contigs with complete CDS, eight incomplete contigs and two pseudogenes were recovered (Fig. S6). Among the TPS contigs with full-length CDS, only the *Meiogyne heteropetala* cineole synthase (*MhCINS*) and the putative TPS2 were recovered in the gene subfamily TPS-b (Fig. S7), comprising largely monoterpene synthases and isoprene synthases of angiosperms (Chen et al., 2011), while the remaining contigs were nested in the subfamily TPS-a, composed of mostly sesquiterpene synthases in angiosperms (Chen et al., 2011). *MhCINS* is one of the top 1% most highly expressed contigs in the transcriptome and has the highest expression level in all 15 TPS contigs detected (Fig. S6; 41–285-fold difference in fpkm).

The transcript of *MhCINS* is 2,076 bp long (accession no.: OQ592534), while the genomic DNA of *MhCINS* is 3,325 bp long (accession no.: OQ592535). The gene consists of five exons and four introns (Fig. S8a). The full-length protein comprises 583 amino acids (Fig. S8b), with the first 39 amino acids at the N-terminal predicted to be the chloroplast transit peptide (probability: 0.96; DeepLoc 2.0; Thumuluri et al., 2022; https://services.healthtech.dtu.dk/services/DeepLoc-2.0/).

### Relative expression level of putative TPS

RT-qPCR revealed the expression pattern of *MhCINS* followed a bimodal distribution (Fig. 5c, d) and appeared unrelated to sex. The non-parametric Kruskal-Wallis test detected a significant difference in floral expression level among putative TPSs (χ^2^ = 18.6, df = 4, *p* = 9.36 × 10^-4^***). In flowers, *MhCINS* has a considerably higher expression level compared to all other TPSs tested (Fig. 5c; *ca*. 70–169-fold difference), congruent with the transcriptome data (Fig. S6). The expression level of *MhCINS* among organs in RT-qPCR was highest in the flowers (Fig. 5d; χ^2^ = 21.6, df = 5, *p* = 0.0006***; *ca.* 2000–25000-fold difference). *MhCINS* is a flower-specific TPS, and the only highly expressed TPS in flowers.

### Functional characterisation of MhCINS

The full length MhCINS protein did not show any catalytic activity for GPP. With deletion of the transit peptide, the enzymatic activity of MhCINS was optimized at pH 7.0 and 50 mM MgCl2 (Fig. S9), while enzyme precipitation was observed at all concentrations of MnCl2 tested. No sesquiterpene products were formed after adding FPP. With the addition of GPP, MhCINS produces 21 monoterpene products, with 1,8-cineole (80.45 %) being the major product (see Table 1 for complete product profile). The enzyme has a *Km* value of 9.19 µM, a *Kcat* value of 2.65 × 10^-2^ min^-1^, and a *Kcat/Km* value of 2.8 × 10^-3^ min^-1^ µM^-1^, comparable to the cineole synthase of *Arabidopsis thaliana* (Chen et al., 2004). The floral terpene profile lacks linalool and has a much lower content of α-terpineol than the MhCINS product profile, likely owing to the difference in headspace-based extraction method of floral odour and solvent-based extraction method of enzyme products, which would presumably lead to underrepresentation of molecules with lower vapour pressure in the floral odour (Table S2).

### Bioassay

We detected pervasive singularity issues in all GLMM models, which was only resolved after all mixed terms were removed. Therefore, we presented the results of binomial tests (Fig. 5e). In the two-choice bioassays, *L. sharpi* was shown to prefer the odour of *M. heteropetala* flowers (72.7%; *p=*0.0262*) and aerial litter (78.2%; *p*=0.0053**) over the air control. The beetles preferred both flower synthetic mix (74%; *p*=0.0173*) and MhCINS synthetic mix (80%; *p*=0.002**) over the solvent controls. *Loberus sharpi* was attracted to the major floral molecule 1,8-cineole over the solvent control (78.6%; *p*=0.0019**). Furthermore, the pollinator did not differentiate between the floral blend and MhCINS blend (*p*=0.29, ns), suggesting MhCINS alone is sufficient to explain olfactory attraction. When subjected to both the floral blend and 1,8-cineole, *L. sharpi* showed no preference (*p*=0.5, ns), possibly indicating a broad preference for varying volatile blends. Lastly, the beetles were attracted to aerial leaf litter over flowers (72.0%, *p*=0.0216*). Unfortunately, we did not collect the odour blends from the individual leaf litter samples which the beetles prefer over floral tissues, so we cannot determine the compositional blends from these litter samples for comparison with the known blends from other assays.

## Discussion

### Cognitive misclassification

Floral visitors of *M. heteropetala* consisted of arachnids, blattodeans, coleopterans and katydids, which are commonly associated with aerial litter (Gradwohl & Greenberg, 1982; Greenberg, 1987; Nadkarni & Longino, 1990; Rosenberg, 1993; Rosenberg, 1997; Snaddon et al., 2012; Mansor et al., 2019). The erotylid beetle *Loberus sharpi* is the most likely effective pollinator of *M. heteropetala*. The family Erotylidae are known to have diverse dietary specialisations, including mycophagy, phytophagy and saprophagy (Carlton et al., 2000; van Zandt et al., 2003; Robertson et al., 2004). Our study revealed that *L. sharpi* specialised in aerial litter because adults and larvae were found exclusively in this substrate in our field sites, except when they were found in *M. heteropetala* flowers. As is typical for the Annonaceae, the perianth of *M. heteropetala* flowers abscise and drop to the forest floor immediately after anthesis, where the larvae do not persist, meaning that eggs laid within floral tissues are very unlikely to reach adulthood (a typical outcome for floral mimicry of brood site substrates). It is possible that abscised floral tissues could become trapped as aerial litter, suggesting that the beetles’ oviposition in flowers represents early colonization of a true brood substrate. However, we failed to find beetle oviposition taking place in flowers of any co-occurring species, nor did we see oviposition occurring on other living vegetative tissues. Collectively, these observations suggest the beetles utilise leaf litter once it has been trapped within the canopy, and that oviposition in *M. heteropetala* flowers is a misguided behaviour resulting from cognitive misclassification of the flowers as aerial litter, with the flowers tapping into a pre-existing relationship between the beetles and their litter substrates.

### Visual and olfactory advergence

The visual cues of *M. heteropetala* flowers are highly similar to aerial litter. The spectral reflectance of flowers matched a subset of aerial litter and was nested within its larger range of spectra, suggesting the flower and litter substrates matches strongly in colour (Fig. 3c). Likewise, the flowers fall within the sizes of aerial litter that the beetles would encounter, though their medians differ. High variation in colour and size of aerial litter is likely due to heterogeneity in leaves in the community and the random quantity of litter per cluster.

Similarly, *ca.* 99% floral volatiles (relative abundance) were also found in aerial litter, although the volatile composition of aerial litter is quantitatively and qualitatively highly variable. The high variation in litter volatiles is expected from the heterogeneity and random assortment of litter (also microclimate variation, age of litter, etc.), so it is not surprising that the floral samples do not overlap strongly with the litter samples on the NMDS (Figure 4b). More importantly, the proximity of floral scent samples to a number of the litter odour samples, combined with the relatively larger distance between *Meiogyne heteropetala* and three closely related congeners suggest olfactory advergence towards volatile blends typical of aerial litter. Together, advergence of floral cues and misguided oviposition behaviour of the beetles provide unequivocal evidence for floral mimicry of aerial litter in *M. heteropetala*. As far as we are aware, this is the first report of aerial litter mimicry.

### Imperfect floral mimicry of aerial litter

Collectively, our data show that the floral phenotype of *Meiogyne heteropetala* strongly resembles a subset of the aerial litter in the visual and olfactory spaces. Although the size, colour and odour of aerial litter are broad and heterogeneous, it is likely that a narrower range of cues indicate to the pollinator which clumps of aerial litter are best suited for oviposition. Current knowledge of the sensory perception and preferences for insects utilising this ecological niche is limited. Further study of these insects is necessary to better determine whether the phenotype of mimetic flowers closely matches the cues of the optimal brood sites. Furthermore, imperfect mimicry could arise, for example, when multiple preferred models are involved, in which pollinators rely on more general cues and therefore exert loose selection over the mimics’ phenotypes (Edmunds, 2000). *Loberus sharpi* has likely developed flexible odour preferences to accommodate the highly variable origin of aerial litter (Fig. 4b; Table S3). Overall, the presence of *Loberus sharpi* and other litter-associated insects at flowers of *M. heteropetala* strongly suggests that the floral phenotype overlaps sufficiently with cues used by these insects to locate aerial litter substrates.

The only result that potentially draw questions on the effectiveness of *M. heteropetala* as an aerial litter mimic is our bioassay result showing that *L. sharpi* adults were significantly more attracted to the scent of aerial litter over that of the flowers. This result contrasts with other bioassay results that unequivocally showed that beetles significantly preferred floral scents over empty controls, and the field observation of oviposition which clearly indicates cognitive misclassification. We did not sample the odour blend from the aerial litter used in the bioassays, so we are unable to determine what chemical constituents contribute to their difference; however, this curious result does suggest that there may be a broader range of acceptable and preferred substrate cues sought by the beetles. *Meiogyne heteropetala* probably only exploited general search cues that indicate general but not the most preferred substrates. The plant can still presumably attain reproductive success if the floral cues are sufficiently attractive to draw a necessary threshold of active pollinators.

### Quasi-Batesian mimicry of oviposition site

Establishing evidence for floral mimicry (vs. systems involving honest floral resources) can be challenging when the pollinator obtains some level of rewards (i.e. quasi-Batesian mimicry *sensu* Speed, 1999). For example, in the *Dracula* orchids that mimic mushrooms, mycetophagous flies still obtain food and shelter on the flowers as they would on the fungal models, but the larvae laid on flowers inevitably perish (Policha et al., 2016, 2019). The current study shows that *M. heteropetala* offers honest shelter and tryst site for *L. sharpi*. In particular, the floral size filter (Fig. 1e) strongly limited the access of potential predators. The flowers nonetheless appeared to be an inferior brood site. Given the understorey habitat of *M. heteropetala* and pendent orientation of the flowers, most abscised petals fall directly onto the forest floor where the larvae do not thrive. Larval mortality may be attributed to increased predation risks, pathogen stress and unfavorable changes in microclimate and decomposer community associated with the forest floor (Moore & Stephenson, 2003; Marques et al., 2015). However, *ca.* 80% of aerial litter was secured to branches in some ways (Fig. S1). This suggests that pre-existing aerial litter dominated by anchored litter provides a much safer brood resource. While the flower offers honest shelter and tryst site rewards, the risky oviposition-site reward is suboptimal, suggesting *M. heteropetala* is a quasi-Batesian mimic.

### Biochemical basis for chemical mimicry

Efforts to characterise TPSs in Annonaceae is largely restricted to Ylang Ylang (*Cananga odorata* var. *fruticosa*), in which one monoterpene synthase and four sesquiterpene synthases have been implicated in the production of floral fragrance (Jin et al., 2015; Dhandapani et al., 2020). Olfactory mimicry of aerial litter in *M. heteropetala*, however, was largely enabled by the monoterpene synthase MhCINS. In plants, terpenoids are synthesised by TPSs (Tholl, 2006; Chen et al., 2011; Karunanithi & Zerbe, 2019). In *M. heteropetala*, the flower-specific MhCINS was responsible for olfactory mimicry. Because the catalytic process of TPSs involves the formation of highly reactive carbocation intermediates, which are subject to a cascade of various events such as isomerisation, cyclisation, hydride shift and water capture (Christianson, 2006; Tantillo, 2011), TPSs produce a diversity of terpenoid products. For the same reason, over half of monoterpene and sesquiterpene synthases in plants can generate multiple products from a single substrate (Degenhardt et al., 2009). Product profile is determined by the environment of the catalytic sites of the TPSs (Christianson, 2006), and has a biochemical constraint. The terpene products are therefore linked and represent a single unit of selection. With less stringent pollinator preference as discussed above, TPSs could probably afford to produce monoterpene blends with components and ratios which may otherwise be unfavorable.

The combinations of floral cues can be regarded as peaks in an adaptive landscape for pollination fitness (Wright, 1932; Simpson, 1944, 1953; Raguso, 2003). Relatively few olfactory components appear to constitute the adaptive peaks of aerial litter mimics, or at least, are required to elicit pollinator attraction. Likewise, in other oviposition-site mimics, only a few key volatiles, such as skatole and indole in the dung mimic *Wurmbea elatior* (Johnson et al., 2020) and oligosulphides in the carrion mimics *Eucomis bicolor* and *Eucomis humilis* (Shuttleworth & Johnson, 2010), are sufficient to at least attract pollinators: the addition of these compounds to non-deceptive congeneric species can shift pollinator makeup to almost entirely saprophilous insects, although other non-olfactory traits also appeared to play a role in pollinator attraction. Moreover, the phenotypic changes associated with new adaptive peaks could happen in a few generations (Gervasi & Schiestl, 2017). The simple biochemical basis for olfactory mimicry of aerial litter suggests that it may be much more prevalent than the literature suggests.

### Potential defence role of floral volatiles

Pollination systems involving 1,8-cineole as the primary floral volatile appear to be context-specific and restricted to a small number of species, such as moth-pollinated *Nicotiana* (Raguso et al., 2003), Euglossine bee-pollinated *Catasetum* (Hills et al., 1972), and *Annona glabra* with specialisation in Chrysomelidae pollinators (Gottsberger, 1999; Goodrich & Raguso, 2009). Nonetheless, 1,8- cineole has been reported in numerous lineages as a secondary floral scent component (Knusden et al., 2006). Monoterpenes are thought to correlate with the repellence of herbivores and facultative floral visitors (Junker & Blüthgen, 2010; Schiestl, 2010). Their roles as toxins and repellents for herbivores, parasites and microbes have been extensively studied (Tholl, 2006; Chen et al., 2011), including the roles of 1,8-cineole (Lee et al., 2004; Dhakad et al., 2018). In several *Eucalyptus* species, 1,8-cineole was shown to effectively reduce the defoliation and dieback of trees caused by Christmas beetles (*Anoplognathus*) (Edwards et al., 1993). It would not be surprising if 1,8-cineole could act as both pollinator attractant and antagonist deterrent in *M. heteropetala*, as such dual functions indeed exist for some terpenoids (Zhou et al., 2017; Pichersky & Raguso, 2018). However, given the floral specificity of MhCINS, the hypothetical antagonists likely represent specialised florivores rather than general herbivores. The elevated floral content of 1,8-cineole might be the result of concerted selective pressure from pollinator attraction and defence against floral antagonists, and may not necessarily be optimised for pollinator attraction, thereby offering an alternative explanation for imperfect olfactory mimicry. Interplays between pollinator attraction and defence in floral chemistry have been demonstrated to adversely affect pollinator visitation but presumably improve overall fitness in other systems (Galen et al., 2011; Ramos & Schiestl, 2019). 1,8-cineole could be initially selected for defence against floral antagonists, serving as an evolutionary step towards olfactory mimicry of aerial litter. A similar hypothesis was invoked to account for the evolutionary origin of aphid mimicry in *Epipactis*, postulating that monoterpenes initially attract non-pollinating predators to keep aphids at bay (Stökl et al., 2011). Future investigation on the potential defensive role of MhCINS may shed light on the evolutionary origin of aerial litter mimicry.

### Implication of low visitation rate

Visitation by *L. sharpi* was strikingly low in this system. Low visitation rate however, is common in some deceptive systems, such as generalised food deceptive (Jersáková & Johnson, 2006) and sexual mimetic systems (Whitehead & Peakall, 2009). Pollinators in these systems adopt longer flight distance, in turn promoting gene flow and genetic diversity (Johnson & Schiestl, 2016 and references therein). It is unclear whether aerial litter mimicry would likewise promote xenogamy. *Meiogyne heteropetala* may offer an invaluable system for studying the influence of specialised pollination strategy on genetic diversity.

### Other floral adaptation and future studies

Non-olfactory floral adaptation is evident in *M. heteropetala*. The most striking specialisation perhaps is dark corolla, which has probably evolved only once in *Meiogyne.* There are only two closely related species which produce maroon or indigo petals: *Meiogyne bidwillii* on the east coast of Queensland and an undescribed species from Groote Eylandt, Australia. Based on our unpublished data, the former predominantly emits an ether volatile whereas the undescribed species emits a minty floral odour like *M. heteropetala* and may potentially represent another aerial litter mimic. Future study of *M. heteropetala* and related species using a combination of anthocyanin characterisation, differential gene expression analysis based on transcriptomic data, and visual bioassay could help elucidate the importance of visual mimicry for aerial litter mimics. Furthermore, oviposition was restricted to the inner petal corrugation (Fig. 2c), suggesting the warty structure (Fig. 1f) may provide an important component for potential tactile mimicry. The inner petal corrugation has previously been regarded as a gland despite the absence of secretary openings (Xue et al. 2021); its actual function has yet to be elucidated. Because the Annonaceae lacks specialisation in pollen presentation, tactile cues like the petal corrugation could encourage the beetles to linger in the flowers, thereby increasing pollen load. Interestingly, dark warty floral structures have evolved independently in *Chiloglottis*, and are believed to offer tactile cues in sexual deception (Wong et al., 2017b; 2022). Apart from potential tactile cues, other floral specialisations, such as size filter, orientation, and floral architecture are exciting avenues to study floral mimicy.

## Conclusions

Our findings show that *M. heteropetala* produces visual and olfactory cues similar to aerial litter, deceiving the aerial litter specialist *Loberus sharpi* for pollination. The current study expands our knowledge of the ecological and biochemical basis of floral mimicry in early-divergent angiosperms and represents the first reported case of aerial litter mimicry. Due to the ubiquity of aerial litter in tropical and temperate forests and its associated arthropod communities, this study paves the way for understanding the interactions between plants and aerial litter arthropods. Our study also contributes to the delineation of the biochemical basis of deceptive pollination systems.

## Acknowledgements

We would like to thank Alison and Steve Pearson for field assistance, Garry and Nada Sankowsky, Janelle Jung and Gardens by the Bay, Aaron Bean and Mackay Regional Botanical Garden, for access of their living collections, Dr Vivian Sandoval Gómez and Queensland Museum for assisting beetle identification, and Jessie Lai and Laura Wong for technical support. We would also like to thank Dr Robert A. Raguso for comments on the manuscript. This research is funded by the Hong Kong Research Grants Council (HKU17112616), awarded to R.M.K.S.

## Author Contributions

M.F.L. and R.M.K.S. planned the project. M.F.L., J.C., P.C.C. and T.S. conducted the field work. M.F.L. and D.C. designed the lab experiments. M.F.L. conducted the lab experiments, performed the analysis and wrote the manuscript. M.F.L., J.C.,

K.R.G. and R.M.K.S. provided interpretation.

## Data Availability

Sequences generated are deposited on National Centre for Biotechnology Information (NCBI).

## Supporting Information

**Fig. S1** Categorisation of aerial litter based on anchorage types in arboreal environments.

**Fig. S2** Principal component analysis of spatial dimension of *Meiogyne heteropetala* flowers and co-occurring aerial litter clumps.

**Fig. S3** Variation in floral scent composition among populations and sexual phases.

**Fig. S4** Kyoto Encyclopedia of Genes and Genomes (KEGG) in the floral transcriptome of *Meiogyne heteropetala*.

**Fig. S5** Anthocyanin and flavonol biosynthesis in *Meiogyne heteropetala*

pistillate-phase flowers.

**Fig. S6** Terpenoid biosynthesis in *Meiogyne heteropetala* pistillate-phase flowers.

**Fig. S7** Maximum likelihood tree of 75 terpene synthase protein sequences. **Fig. S8** Genomic DNA sequence of *MhCINS* and alignment of related TPSs. **Fig. S9** MhCINS activities at varying pH values and MgCl2 concentrations. **Fig. S10** Variance Inflation Factor (VIF) of bioassay responses.

**Table S1** List of primers used in the study.

**Table S2** List of synthetic terpenoids used in solvent mix for the two-choice bioassay.

**Table S3** Volatile matrix of MhCINS, *Meiogyne heteropetala* organs, and aerial litter.

**Table S4** SIMPER analysis (flower vs aerial litter).

**Methods S1** Supplementary Methodology.

