## Supplemental Data 1 for "Aerial litter mimicry: a novel form of floral deception mediated by a monoterpene synthase"

The following Supporting Information is available for this article:

**Fig. S1** Categorisation of aerial litter based on anchorage types in arboreal environments.

**Fig. S2** Principal component analysis of spatial dimension of *Meiogyne heteropetala* flowers and co-occurring aerial litter clumps.

**Fig. S3** Variation in floral scent composition among populations and sexual phases.

**Fig. S4** Kyoto Encyclopedia of Genes and Genomes (KEGG) in the floral transcriptome of *Meiogyne heteropetala*.

**Fig. S5** Anthocyanin and flavonol biosynthesis in *Meiogyne heteropetala* pistillate-phase flowers.

**Fig. S6** Terpenoid biosynthesis in *Meiogyne heteropetala* pistillate-phase flowers.

**Fig. S7** Maximum likelihood tree of 75 terpene synthase protein sequences.

**Fig. S8** Genomic DNA sequence of *MhCINS* and alignment of related TPSs.

**Fig. S9** MhCINS activities at varying pH values and MgCl2 concentrations.

**Fig. S10** Variance Inflation Factor (VIF) of bioassay responses.

**Table S1** List of primers used in the study.

**Table S2** List of synthetic terpenoids used in solvent mix for the two-choice bioassay.

**Methods S1** Supplementary Methodology.

**Fig. S1** Categorisation of aerial litter based on anchorage types in arboreal environments. (a) Aerial litter self-retained by vasculature, derived from dried dead plant tissues that did not abscise from the plant. (b) Aerial litter anchored by arthropod silk, derived from abandoned nests of green tree ants. (c) Aerial litter anchored by arthropod silk (indicated by yellow arrows), derived from brood shelters utilised by spiders. (d) Aerial litter anchored by marasmioid fungal rhizomorph network (indicated by blue arrows), which traps falling litter from above. (e) Free-hanging aerial leaf litter. (f) Composition of aerial litter based on its anchorage type (Dingo beach: *n*=50; Conway National Park: *n*=38). (Distinction for the type of arthropod silk was not made here due to assessment difficulty. Green tree ant nests that were actively in use was also omitted because their foliage was often green.)

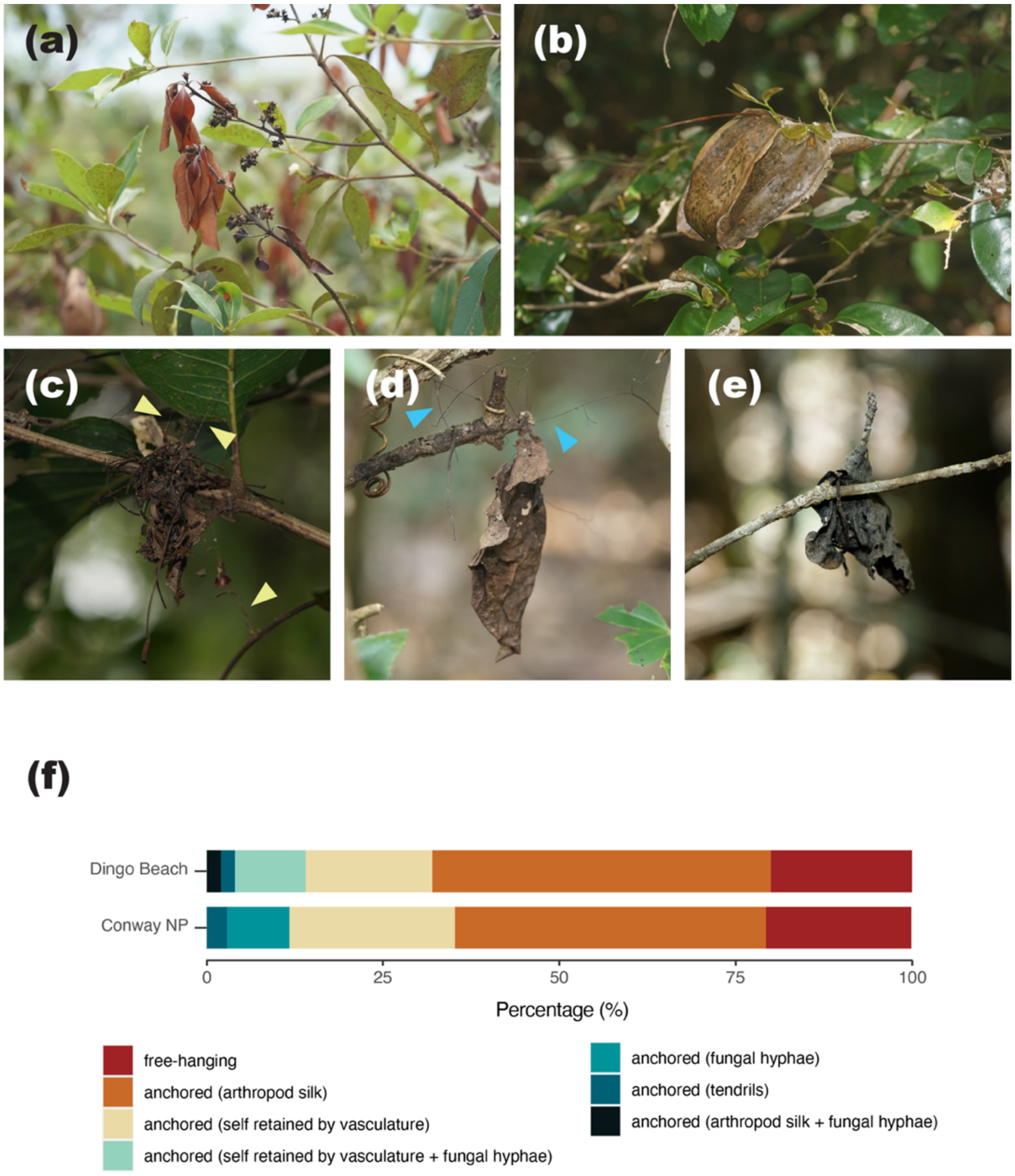

**Fig. S2** Principal component analysis of the spatial dimension of *Meiogyne heteropetala* flowers and co-occurring aerial litter clumps.

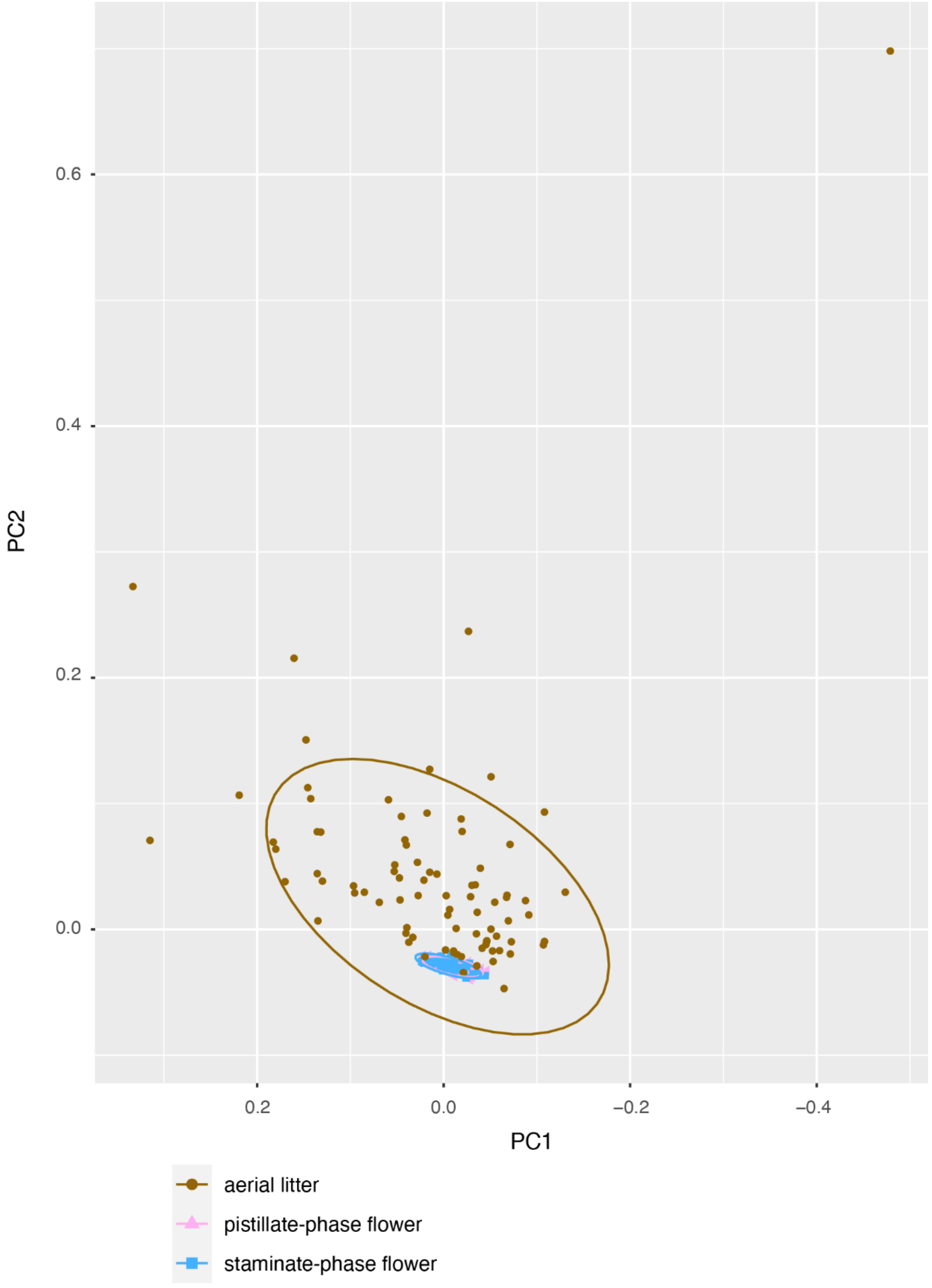

**Fig. S3** Variation in floral scent composition among populations and sexual phases determined using dynamic headspace extraction method (total $n$=24; NMDS based on Bray-Curtis dissimilarity, 2D stress = 0.1491).

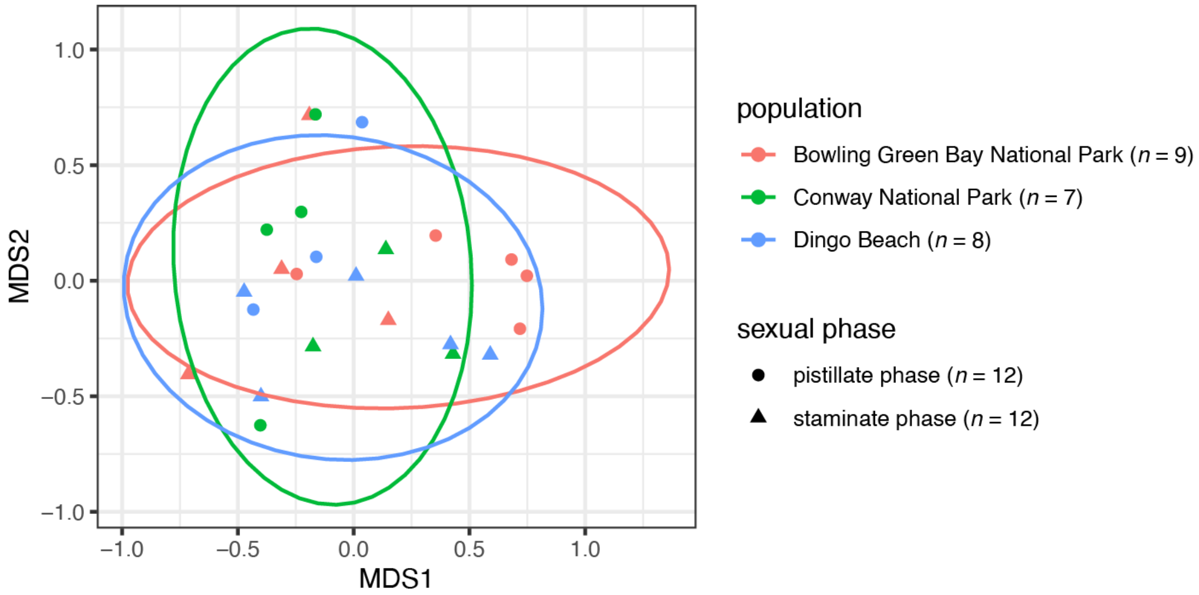

**Fig. S4** Kyoto Encyclopedia of Genes and Genomes (KEGG) annotations of 31,269 contigs identified in the floral transcriptome of *Meiogyne heteropetala*.

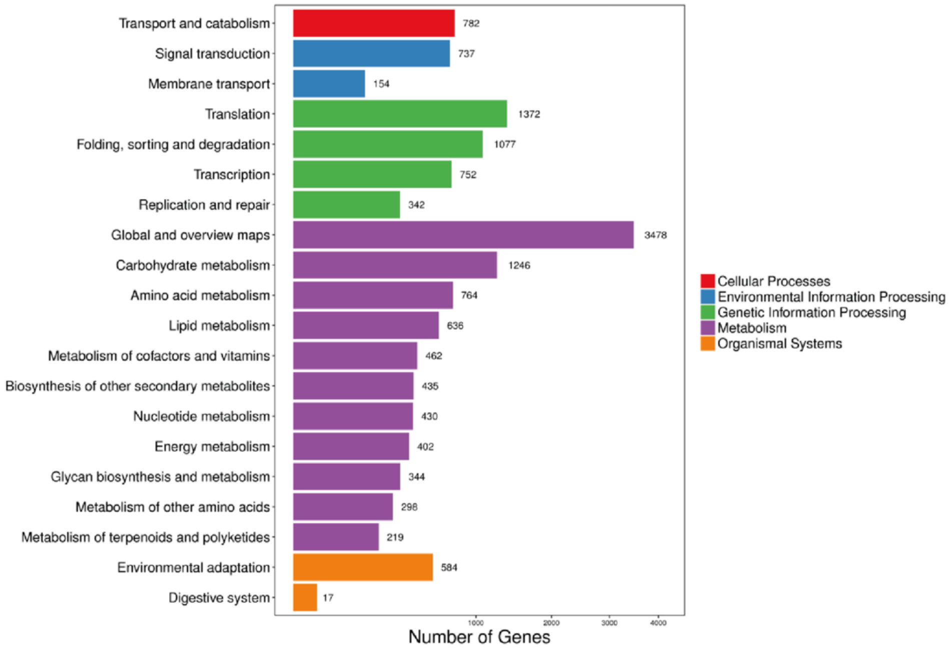

**Fig. S5** Anthocyanin & flavonol biosynthesis in *Meiogyne heteropetala* pistillate-phase flowers. Expression level of transcripts involved in terpenoid biosynthesis in the floral transcriptome are denoted by $\log_2(\text{fpkm}+1)$ value (fpkm: fragments per kilobase of transcript per million reads mapped). Transcripts with top 1% expression level in the whole transcriptome are indicated by red boxes. 3AT: anthocyanidin 3-O-glucoside 6″-O-acyltransferase; ANR: anthocyanidin reductase; ANS: anthocyanidin synthase; C3′H: 5-O-(4-coumaroyl)-D-quinate 3′-monooxygenase; BZ1: anthocyanidin 3-O-glucosyltransferase; C4H: cinnamate 4-hydroxylase; CCoAOMT: caffeoyl-CoA O-methyltransferase; CHI: chalcone isomerase; CHS: chalcone synthase; F3H: flavanone 3-hydroxylase; F3′H: flavonoid 3′-hydroxylase; FLS: flavonol synthase; GT1: anthocyanidin 5,3-O-glucosyltransferase; HCT: shikimate O-hydroxycinnamoyltransferase; DFR: dihydroflavonol 4-reductase; UGT79B1: anthocyanin 3-o-glucoside 2″-o-xylosyltransferase.

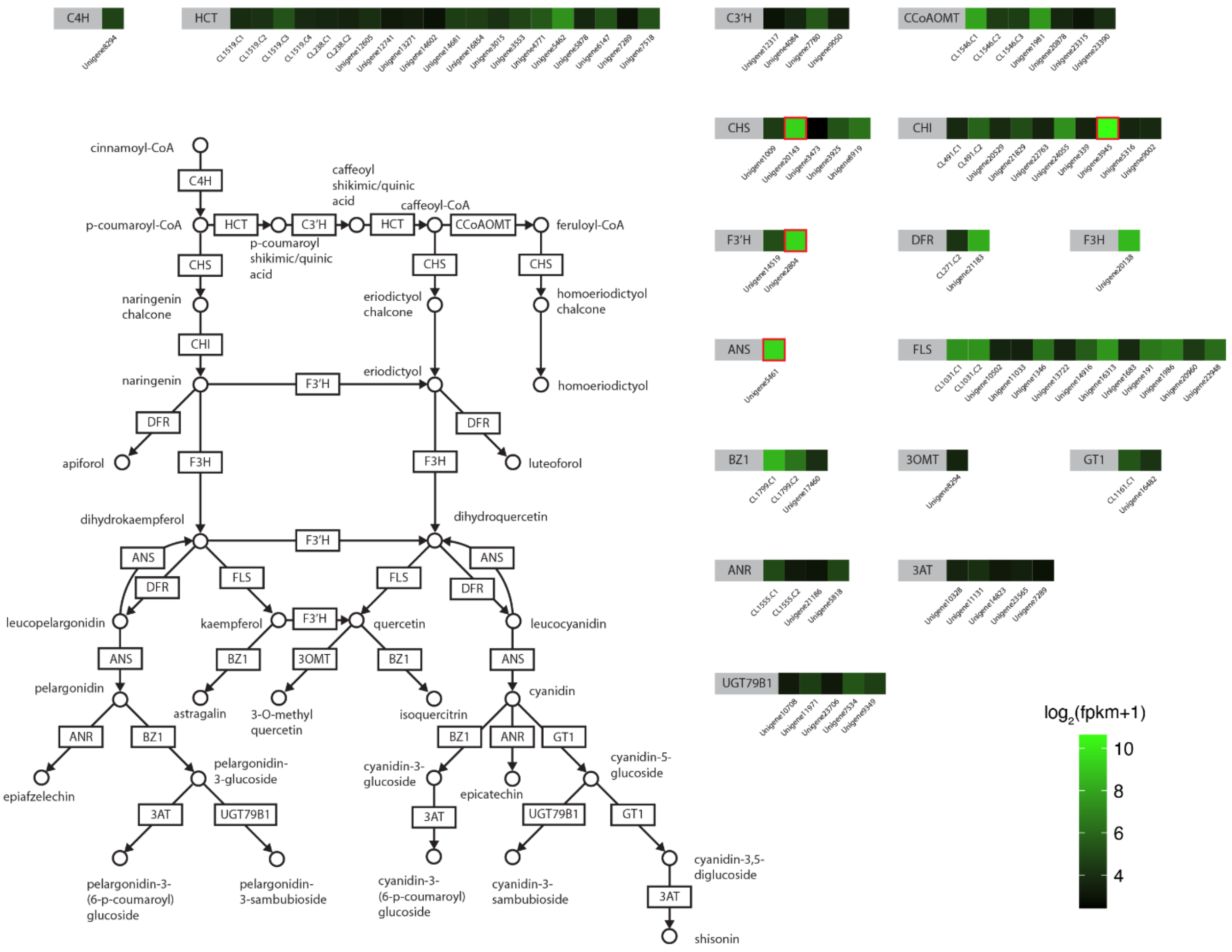

**Fig. S6** Terpenoid biosynthesis in *Meiogyne heteropetala* pistillate-phase flowers. Expression level of enzymes involved in terpenoid biosynthesis in the floral transcriptome are denoted by log2(fpkm+1) value (fpkm: fragments per kilobase of transcript per million reads mapped). Transcripts with top 1% expression level in the whole transcriptome are indicated by red boxes. ACAT: acetoacetyl-CoA thiolase; CDP-ME: 4-diphosphocytidyl-2-C-methylerythritol; CDP-MEP: 4-diphosphocytidyl-2-C-methyl-D-erythritol 2-phosphate; DMAPP: Dimethylallyl pyrophosphate (DMAPP); DXP: 1-Deoxy-D-xylulose 5-phosphate; dxr: DXP reductoisomerase; dxs: DOXP synthase; FDPS: farnesyl diphosphate synthase; GDPS: geranyl diphosphate synthase; HMB-PP: (E)-4-Hydroxy-3-methyl-but-2-enyl pyrophosphate; HMGCS: HMG-CoA synthase; HMGCR: HMG-CoA reductase; IDI: isopentenyl pyrophosphate isomerase; IPP: Isopentenyl pyrophosphate; IspD: 2-C-methyl-D-erythritol 4-phosphate cytidylyltransferase; IspE: 4-diphosphocytidyl-2-C-methyl-D-erythritol kinase; IspF: 2-C-methyl-D-erythritol 2,4-cyclodiphosphate synthase; IspG: HMB-PP; IspH: HMB-PP reductase; MEcPP: 2-C-methyl-D-erythritol 2,4-cyclodiphosphate; MEP: 2-C-methylerythritol 4-phosphate; MvaK2: phosphomevalonate kinase; MVD: mevalonate-5-pyrophosphate decarboxylase; MVK: mevalonate-5-kinase; TPS: terpene synthase.

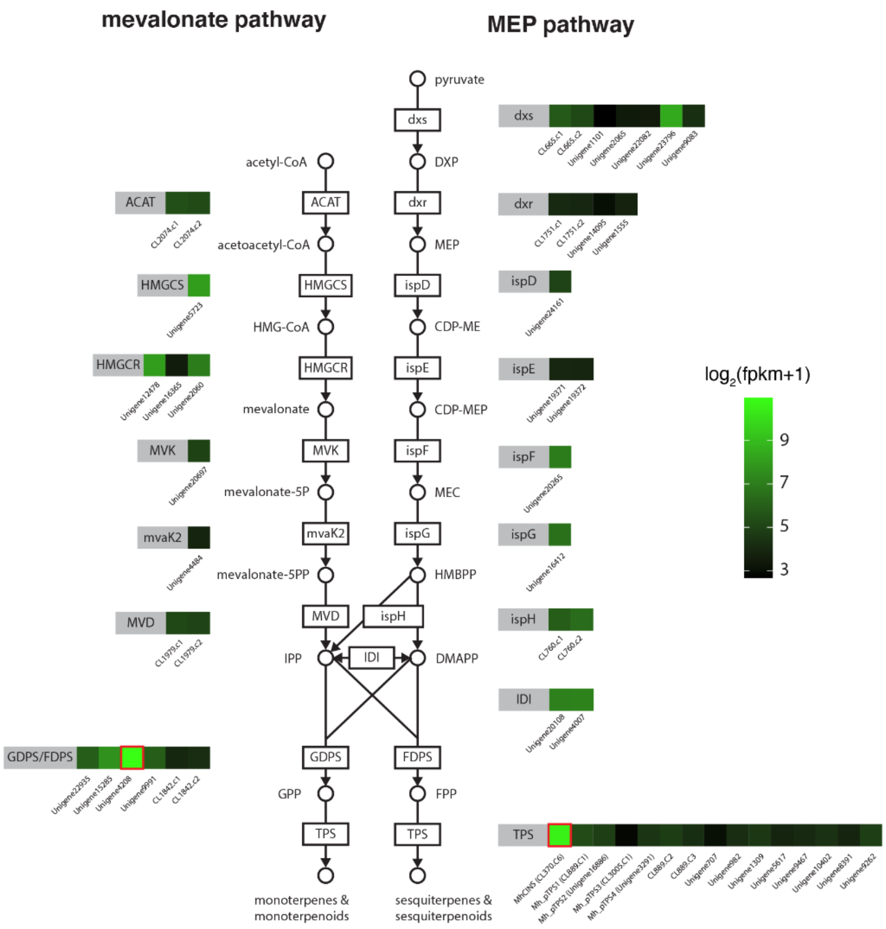
mevalonate pathway
MEP pathway
pyruvate
dxs
DXP
dxr
MEP
ispD
CDP-ME
ispE
CDP-MEP
ispF
MEC
ispG
HMBPP
ispH
DMAPP
IDI
FDPS
FPP
TPS
sesquiterpenes & sesquiterpenoids
acetyl-CoA
ACAT
acetoacetyl-CoA
HMGCS
HMG-CoA
HMGCR
mevalonate
MVK
mevalonate-5P
mvaK2
mevalonate-5PP
MVD
IPP
GDPS
GPP
monoterpenes & monoterpenoids
GDPS/FDPS
log2(fpkm+1)
9
7
5
3

**Fig. S7** Maximum likelihood tree of 75 terpene synthase protein sequences with 1,000 non-parametric bootstraps under the best-fitting Jones-Taylor-Thornton (JTT) substitution model for amino acid from RAxML 8. Numbers on the nodes refer to the bootstrap values. Scale bar denotes substitution rate per site.

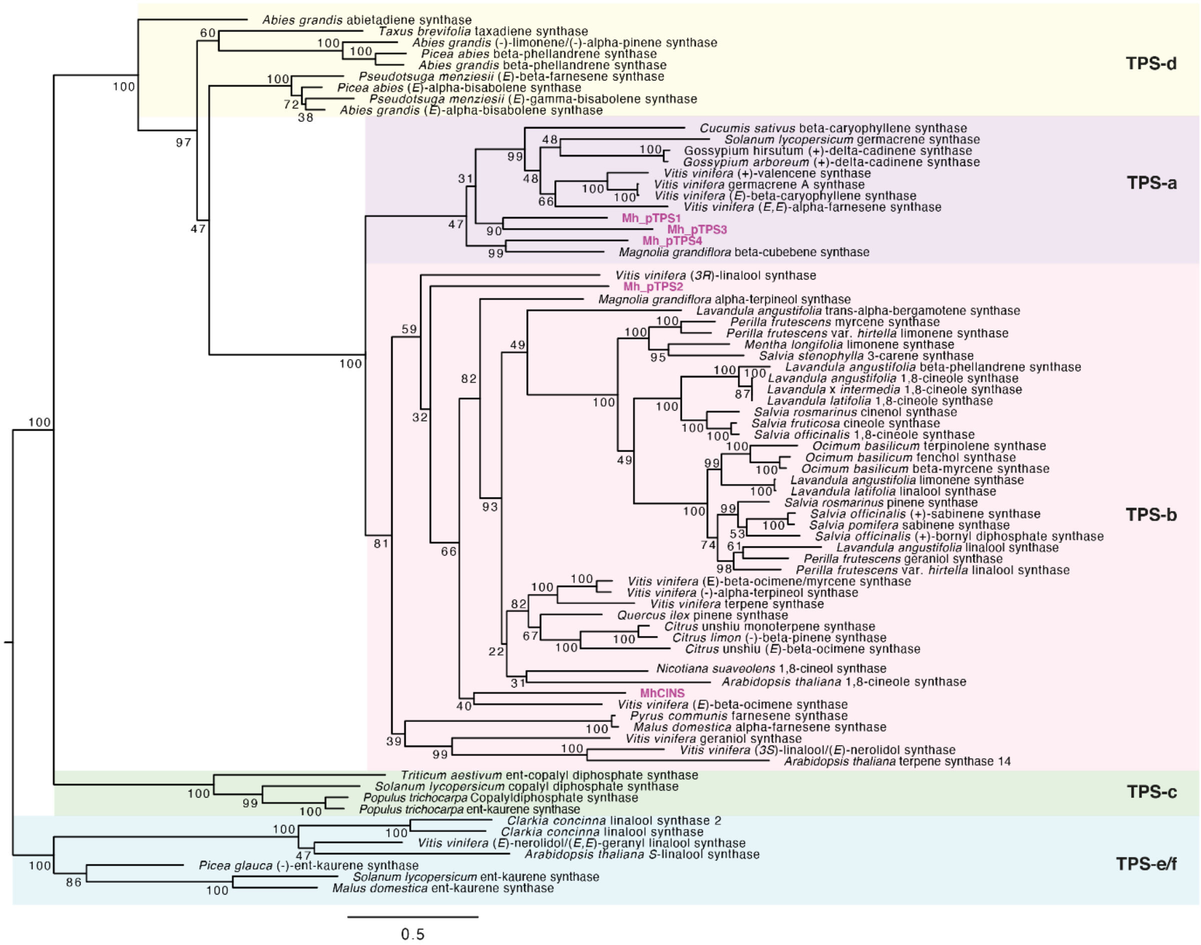

**Fig. S8** Genomic DNA sequence of *MhCINS* and alignment of related TPSs. (a) Schematic representation of *MhCINS* genomic DNA. Exons (1–5) are denoted by rectangular boxes, while introns (1–4) are denoted by interleaving lines. (b) Clustal Omega alignment of the amino acid sequences of MhCINS and five other published monoterpene synthases. Shades indicate the level of sequence similarity. Motifs are annotated by red boxes. The amino acid sequence of the transit peptide of MhCINS is underscored (predicted in DeepLoc 2.0 (Thumuluri et al., 2022)). AtCINS: *Arabidopsis thaliana* cineole synthase (NP_189212); E: exon; LaCINS: *Lavandula augustifolia* cineole synthase (JN701461); Mg17: *Magnolia grandiflora* α-terpineol synthase (EU366430); NsCINS: *Nicotiana suaveolens* cineole synthase (ABP88782); SoCINS: *Salvia officinalis* cineole synthase (AAC26016); UTR: untranslated region.

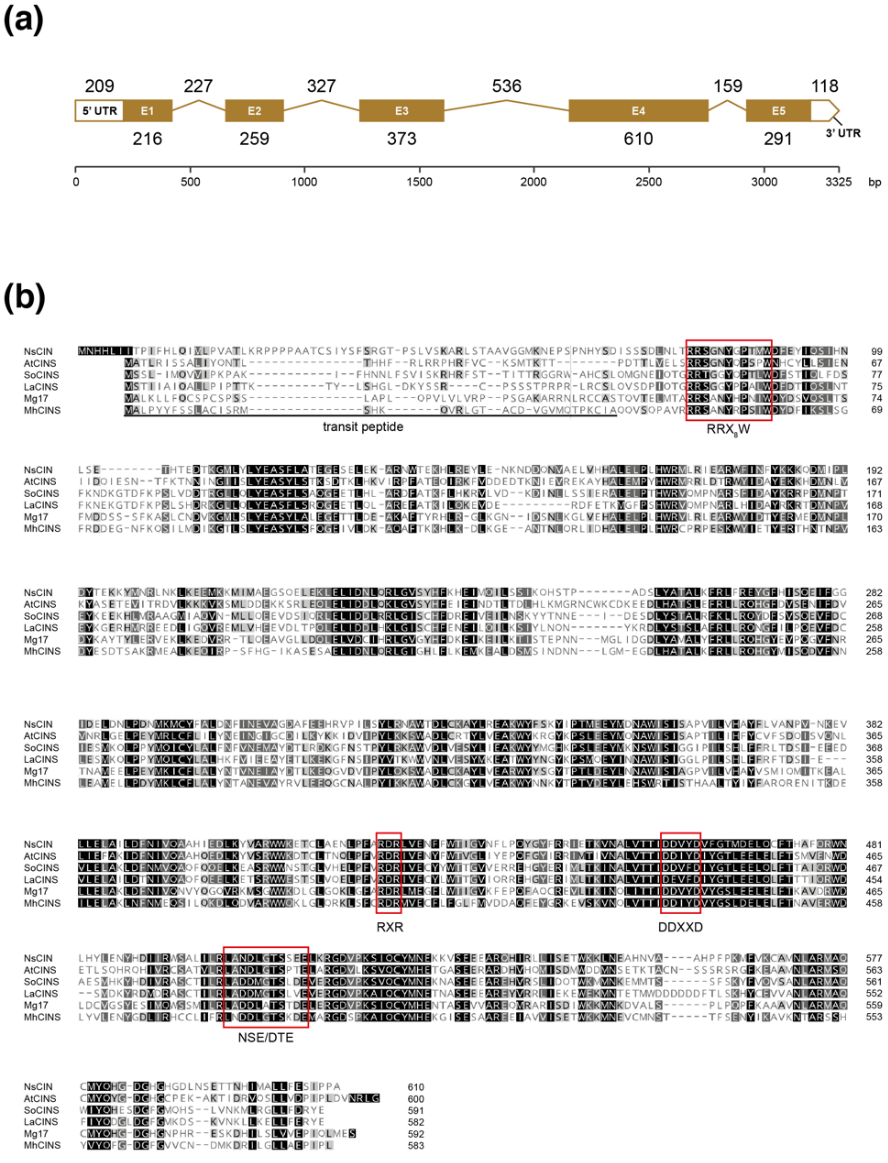

**Fig. S9** MhCINS activities at varying pH values and $MgCl_2$ concentrations. (a) Reaction rates of MhCINS at various pH (pH6.5–9). (b) Reaction rates of MhCINS at various $MgCl_2$ concentration (25–250 mM).

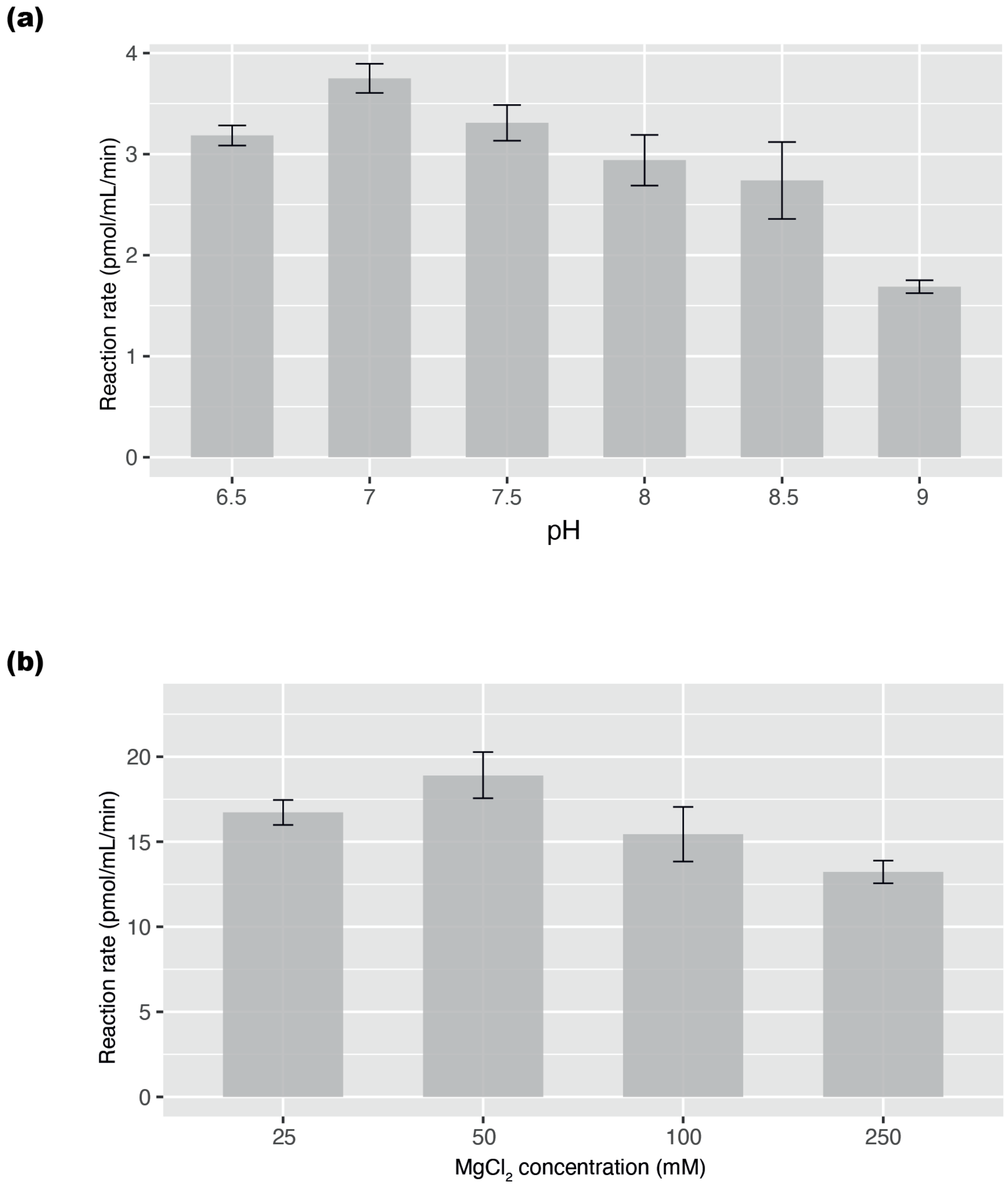

**Fig. S10** Variance Inflation Factor (VIF) of bioassay responses. VIFs from 50 imputed matrices were pooled using the median. Variance in the VIFs were measured using median absolute deviation. (a) VIFs for plant material dataset ($n_{beetles}$=32) (b) VIFs for synthetic mix dataset ($n_{beetles}$=36)

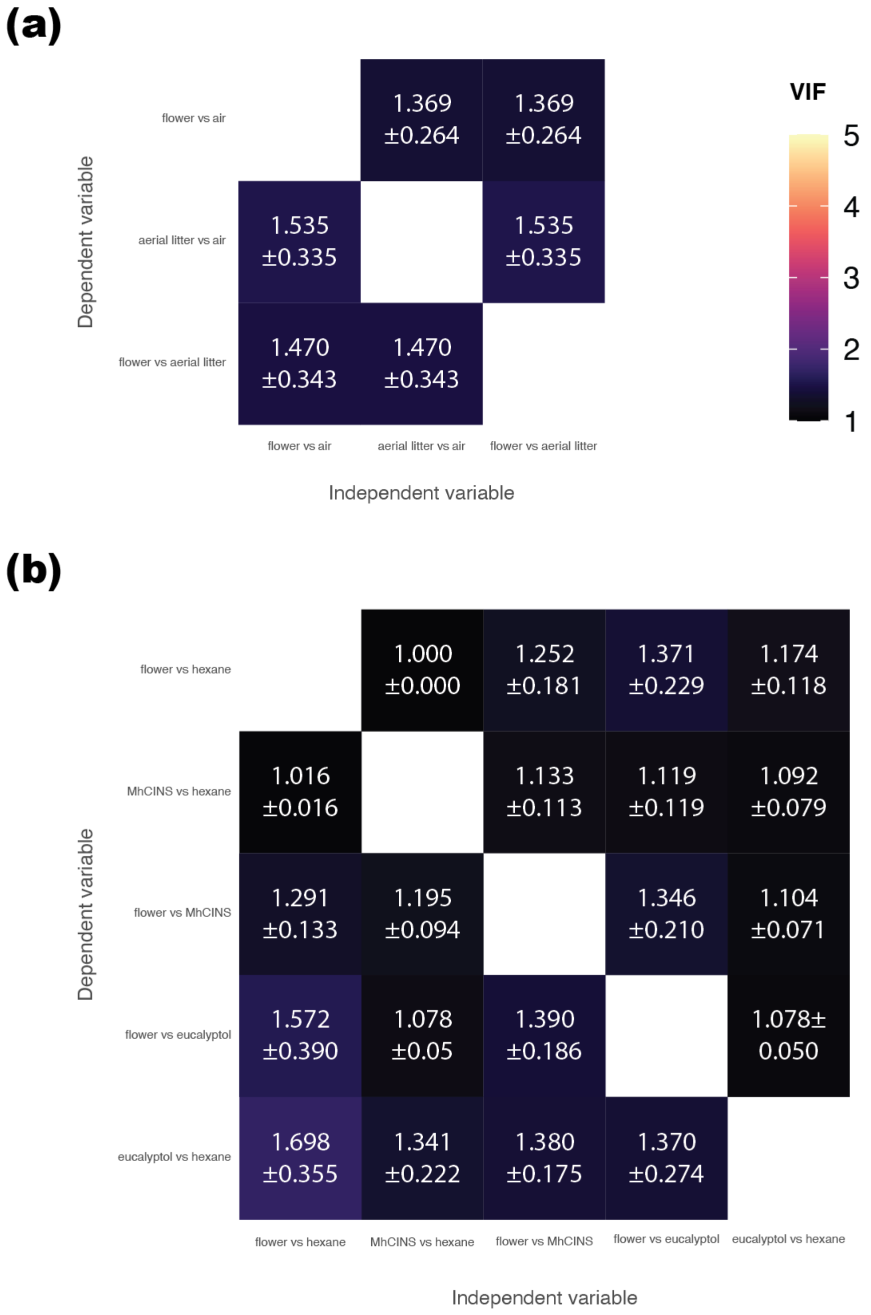

**Table S1** List of primers used in the study.

| Primer Name | Primer sequence (5'–3') |
|---|---|
| MhCINScloning1F | TCTCTATCTAGAAGGAGGTATAATATGGCATTGCCTTACTACT |
| MhCINScloning2F | TCTCTATCTAGAAGGAGGTATAATATGCGGAGATCTGCTAATTATC |
| MhCINScloningR | GTGATGGGATCCGCCGTCAAGTGGTATTGGTTCAGC |
| MhCINSsqF | ATGTCCAAAATCTACGTGATAAAT |
| MhCINSsqR | GGCATGACACCTCCATATT |
| MhCINS_sq261F | CATAAACAGGTGCGCTTAG |
| MhCINS_sq307F | AGACTCCAAAATGCATTGC |
| MhCINS_sq854R | CCCTTTTAGATCCTTGAGG |
| MhCINS_sq753F | ACGCTGAGTCTATATGAAGC |
| MhCINS_sq1244R | ATCCAATGAACCATAAACATCAT |
| MhCINS_sq1162F | AATACGGAAGAAAAGAAGTTT |
| MhCINS_sq1647R | CATTTAGCCGGAAAATCAGA |
| MhCINS_sq1556F | ACAACGAGAAAATATAACGAAAG |
| MhCINS_sq1989R | TCATCAACACAACTCACTAGAT |
| MhCINS-qPCRF | CGTATGTCACATAAACAGGTGCG |
| MhCINS-qPCRR | TGCGGACGTATCAGACTCGT |
| ACT2-qPCRF | ACCTGAAGAACACCCAGTGC |
| ACT2-qPCRR | ACCAGAATCCAACACAATACCAGT |
| UBC21-qPCRF | GCTTGTGTGTGATGATTCTAATAT |
| UBC21-qPCRR | GGCAAATCTCACCTGTCTT |
| TPS1-qPCRF | GGATGACATACAATCAAGACAGTTCG |
| TPS1-qPCRR | CCAACGGGACAGCAGACG |
| TPS2-qPCRF | GAGTACCTAGACAATGGGTGG |
| TPS2-qPCRR | CGTTCCATCTCCGCCTTTGA |
| TPS3-qPCRF | AGATGAACTCGAAATCTTTACCAGC |
| TPS3-qPCRR | GACGAGCACCTTTGTTTCTCT |
| TPS4-qPCRF | ATGGATGATATTCAAACAGACAAGGTC |
| TPS4-qPCRR | GCGACGGCATAAGGATAGACt |
| TPS5-qPCRF | CATCATGTCAAACCAGGCGg |
| TPS5-qPCRR | TTTAGAGTACGCTTGATTGCTTCCT |

**Table S2** List of synthetic terpenoids used in solvent mix for two-choice bioassays. Chemicals purchased from Sigma-Aldrich, Inc. *data obtained from PubChem or the good scents company.

| | Commercial product no. | Vapour pressure (mm Hg at 25 °C)* | Flower mix (ng) | MhCINS mix (ng) |
|---|---|---|---|---|
| (-)-α-pinene | P45702 | 4.75 | 0.6 | 0.7 |
| (+)-α-pinene | P45680 | 4.75 | 12.1 | 17.6 |
| β-myrcene | M100005 | 2.09 | 5.4 | 2.6 |
| sabinene | W530597 | 2.6 | 9.1 | 5.9 |
| α-phellandrene | W285609 | 1.86 | 0.3 | 0.0 |
| (-)-β-pinene | 112089 | 2.93 | 0.8 | 0.5 |
| β-Ocimene | W353901 | 1.56 | 0.3 | 0.1 |
| (*S*)-(-)-limonene | 218367 | 1.55 | 10.7 | 2.6 |
| (*R*)-(+)-limonene | 183164 | 1.55 | 8.2 | 2.9 |
| γ-terpinene | 223190 | 1.09 | 2.0 | 0.4 |
| terpinolene | 86485 | 0.74 | 0.6 | 0.1 |
| 1,8-cineole | C80601 | 1.90 | 244.5 | 252.0 |
| (-)-α-terpineol | W304522 | 0.04 | 0.6 | 6.8 |
| (+)-α-terpineol | 83073 | 0.04 | 1.4 | 7.9 |
| (*E*)-β-caryophyllene | W225207 | 0.013 | 3.2 | 0.0 |
| Total | | | 300 | 300 |

**Methods S1** Supplementary methodology.

To measure floral temperature, a digital data-logger (Testo 176-T4, Testo, Titisee-Neustadt, Germany) was connected to two type-K thermocouples (± 0.3°C accuracy), with one inserted into the floral chamber, and the other placed 10 cm away from the flower to measure ambient air temperature. Temperatures were measured every 10 min throughout 5-day anthesis.

Prior to assessment of pollen grains on pistillate-phasae floral visitors with SEM, pollen morphologies of co-occurring flowering species were retreived from literature, which were determined to be distinct from the monosulcate monad of *M. heteropetala* (*Miliusa brahei* (scabrate circular monad; Chaowasku et al., 2008); *Tabernaemontana orientalis* (four-colporate monad; Haberle et al., 2021); *Planchonia careya* (syncolpate monad; Haberle et al., 2021); *Eugenia reinwardtiana* (tricolporate monad; Haberle et al., 2021)).

To assess whether aerial litter offers a safe brood substrate, a 25 m transect was laid in Dingo Beach and Conway National Park, where one randomly selected aerial litter cluster per tree was assessed as either free-hanging or anchored to branches by other agents (e.g., vasculature, silk).

RNA-seq for a pistillate-phase flower was performed on Illumina Hi-seq 4000 sequencing platform at BGI. Total mRNA was captured using the oligo(dT) method before fragmentation. First and second strands of cDNA were synthesised followed by end repair, single adenine addition and adaptor ligation. The cDNA fragments were size-selected by agarose gel electrophoresis and enriched by PCR before sequencing. The Illumina platform generated 108.3 million raw reads (100 bp paired-end) in total. Adaptors, low-quality reads and reads with > 5% unknown bases were filtered and trimmed SOAPnuke (Chen et al., 2018) to generate clean reads. Removal of adaptors and poor reads resulted in 78.9 million clean reads, constituting 72.84% of data generated. *De novo* transcriptome assembly was performed using Trinity v2.0.6 (Grabherr et al., 2011). Contigs with length less than 150 bp were discarded, minimum count for k-mers was set to three and a minimum of three reads were needed to glue two inchworm contigs. Contigs assembled were annotated with NT, NR, GO, KOG, KEGG, SwissProt and InterPro databases using Blastn v2.2.23, Blastx v2.2.23 (Altschul et al., 1990), Diamond v0.8.31 (Buchfink et al., 2015), Blast2GO v2.5.0 (Conesa et al., 2005) and InterProScan5 v5.11-51.0 (Jones et al., 2014). The candidate coding area was predicted using TransDecoder v3.0.1 (https://transdecoder.github.io). The longest ORF was blasted against SwissProt and Pfam databases using Hmmscan (Eddy et al., 2011). KAAS (Moriya et al., 2007) was used to assign KO identifiers and map the contigs to KEGG pathways, allowing identification of contigs involved in anthocyanin and terpene production. Clean reads were mapped to the assembled transcriptome using Bowtie2 v2.2.5 (Langmead & Salzberg, 2012) with the sensitive preset option. Fragments per kilobase of transcript per million reads (FPKM) were calculated using Cufflinks (Trapnell et al., 2010) with default settings.

The homology model of MhCINS was reconstructed using *Citrus sinensis* (+)-limonene synthase (5uv0.1.A) as the template on SWISS-MODEL (Waterhouse et al., 2018). The model was validated with ERRAT (Colovos & Yeates, 1993), VERIFY3D (Eisenberg et al., 1997) and PROCHECK (Laskowski et al., 1993). All residues fall within the allowed region in Ramachandran plot, and 92.7% residues nested within the most favoured region.

To assess the phylogenetic placement of the potential terpene synthases involved in floral scent production, amino acid sequences of five TPSs with full length CDS generated in this study and 70 published TPSs were aligned using Clustal Omega. Phylogenetic relationships of the terpene synthases were reconstructed using maximum likelihood (ML) method performed in RAxML v.8.2.12 (Stamatakis, 2014). PartitionFinder v.2 (Lanfear et al., 2017) was used to deduce the best-fitting amino acid substitution model among all the models available on the programme. A total of 1000 ML trees were inferred from distinct random-addition-sequence Maximum Parsimony starting trees. Non-parametric bootstrapping was subsequently performed with 1,000 iterations. (Accession numbers: *Abies grandis* (-)-limonene/(-)-alpha-pinene synthase:

AF139207; *Abies grandis* abietadiene synthase: AAK83563; *Abies grandis* beta-phellandrene synthase: AF139205; *Abies grandis* (*E*)-alpha-bisabolene synthase: AAC24192 ; *Arabidopsis thaliana* 1,8-cineole synthase: AAU01970/NP_189212; *Arabidopsis thaliana* putative S-linalool synthase: AAL24105; *Arabidopsis thaliana* terpene synthase 14: AEE33874; *Citrus limon* (-)-beta-pinene synthase: AF514288; *Citrus unshiu* (*E*)-beta-ocimene synthase: BAD91046; *Citrus unshiu* monoterpene synthase: BAD91045; *Clarkia breweri* linalool synthase 2: AAD19840; *Clarkia concinna* linalool synthase: AAD19839; *Cucumis sativus* beta-caryophyllene synthase: AAU05952; *Gossypium arboreum* (+)-delta-cadinene synthase isozyme: AAA93064 ; *Gossypium hirsutum* (+)-delta-cadinene synthase: AAC12784; *Lavandula angustifolia* beta-phellandrene synthase: HQ404305; *Lavandula angustifolia* limonene synthase: ABB73044; *Lavandula angustifolia* linalool synthase: ABB73045; *Lavandula angustifolia* trans-alpha-bergamotene synthase: ABB73046; *Lavandula latifolia* 1,8-cineole synthase : JN701460; *Lavandula latifolia* linalool synthase: ABD77417; *Lavandula* x *intermedia* 1,8-cineole synthase: JN701459; *Magnolia grandiflora* alpha-terpineol synthase: EU366430; *Magnolia grandiflora* beta-cubebene synthase: EU366429; *Malus domestica* alpha-farnesene synthase: AAX19772; *Malus domestica* *ent*-kaurene synthase: AFG18184; *Mentha longifolia* limonene synthase: AF175323; *Nicotiana suaveolens* 1,8-cineole synthase: ABP88782; *Ocimum basilicum* beta-myrcene synthase: AAV63791; *Ocimum basilicum* fenchol synthase: AAV63790; *Ocimum basilicum* terpinolene synthase: AAV63792; *Perilla frutescens* geraniol synthase: ABB30218; *Perilla frutescens* myrcene synthase: AAF76186; *Perilla frutescens* var. *hirtella* limonene synthase: AAF65545; *Perilla frutescens* var. *hirtella* linalool synthase: ACN42009; *Picea abies* beta-phellandrene synthase: AAK39127; *Picea abies* (*E*)-alpha-bisabolene synthase: AAS47689; *Picea glauca* (-)-*ent*-kaurene synthase: ACY25275; *Populus trichocarpa* copalyl diphosphate synthase: XP_002306777; *Populus trichocarpa* *ent*-kaurene synthase: EEE81383; *Pseudotsuga menziesii* (*E*)-beta-farnesene synthase: AAX07265; *Pseudotsuga menziesii* (*E*)-gamma-bisabolene synthase: AAX07266; *Pyrus communis* farnesene synthase: AAT70237; *Quercus ilex* pinene synthase: CAK55186; *Salvia fruticosa* cineole synthase: ABH07677; *Salvia officinalis* (+)-bornyl diphosphate synthase: AAC26017

; *Salvia officinalis* (+)-sabinene synthase: AAC26018; *Salvia officinalis* 1,8-cineole synthase: AAC26016 ; *Salvia pomifera* sabinene synthase: ABH07678; *Salvia rosmarinus* cinenole synthase: ABI20515; *Salvia rosmarinus* pinene synthase: ABP01684; *Salvia stenophylla* 3-carene synthase: AF527416; *Solanum lycopersicum* copalyl diphosphate synthase: BAA84918; *Solanum lycopersicum ent*-kaurene synthase: AEP82778; *Solanum lycopersicum* germacrene synthase: AEM05858; *Taxus brevifolia* taxadiene synthase: AAC49310; *Triticum aestivum ent*-copalyl diphosphate synthase: BAH56558; *Vitis vinifera* (-)-alpha-terpineol synthase: AAS79352; *Vitis vinifera* (+)-valencene synthase: AAS66358; *Vitis vinifera* (3*R*)-linalool synthase: ADR74209; *Vitis vinifera* (3*S*)-linalool/(*E*)-nerolidol synthase: ADR74211; *Vitis vinifera* (*E*,*E*)-alpha-farnesene synthase: ADR74198; *Vitis vinifera* (*E*)-beta-caryophyllene synthase: ADR74192; *Vitis vinifera* (*E*)-beta-ocimene synthase: ADR74204; *Vitis vinifera* (*E*)-beta-ocimene/myrcene synthase: ADR74206; *Vitis vinifera* geraniol synthase: ADR74217; *Vitis vinifera* germacrene A synthase: ADR66821; *Vitis vinifera* (*E*)-nerolidol/(*E*,*E*)-geranyl linalool synthase: ADR74219; *Vitis vinifera* terpene synthase: ADR74202)

To account for random effect due to the reuse of beetles across bioassay, binomial generalised linear mixed models (binomial GLMMs) were performed (Mas et al., 2020; Rondoni et al., 2022; Roberts et al., 2023). Missing data were coded when the beetles produced "no choice" results or when the beetles were not subjected to a particular bioassay. Bioassays for plant materials (flower vs. air; aerial litter vs. air; flower vs. aerial litter) and synthetic mixes (the remaining assays) were analysed separately. To reduce missing data, two beetles with all bioassay responses as "no choice" were removed from the plant material dataset (reduced from *n*=34 to *n*=32). Missing data were imputed using MICE (v.3.16.0) package on R (van Buuren & Groothuis-Oudshoorn, 2011), which generated 50 imputed datasets. Binomial GLMM was performed on lme4 (v.1.1-35.1) package on R (Bates et al., 2015), with responses of other bioassays treated as random variables. Results across imputed datasets were pooled in MICE package. Variance inflation factors (VIF) were calculated using car (v.3.1-2) package on R (Fox & Weisberg, 2019). Binomial GLMM invariably produced overfitted singular models for all bioassays, in which the random variables have zero variance. Since singular models have poor power and produce unreliable statistics, we followed the recommendation of the programme developers to stepwise remove problematic random variables from the full mixed models. Singularities were only resolved once all random variables were removed. Singularity can arise when the model is too complex for the data, and given the prevalence of singularity, in this case, it might potentially be because binomial variables contain too little information for our sample size. The rationale for the use of GLMM is that data could be confounded when, for example, beetles with strong preference for 1,8-cineole were disproportionally used across bioassays. Such bias would be reflected on collinearity between responses of different bioassays, i.e. beetles biased to 1,8 cineole over hexane should also be biased to MhCINS over hexane. However, the VIF matrices instead showed weak multicollinearity among responses of different bioassays (Fig. S10; VIFs: 1.000–1.698<5). It is thus unlikely that beetle's preferences were confounded by such dependency. For these reasons, we presented the results of binomial tests in the main text based on the non-imputed datasets.
